# An Oligomeric Lanthipeptide from *Nostoc punctiforme* Promotes Host Association During Early Symbiosis with *Blasia pusilla*

**DOI:** 10.64898/2026.08.14.744816

**Authors:** Nico Brüssow, Daniel Teutsch, Andi Mainz, Roderich D. Süssmuth, Elke Dittmann

## Abstract

Nitrogen-fixing *Nostoc* species form symbiotic relationships with diverse plants, yet the role of specialized metabolites in these interactions remains poorly understood. Here, we identify a previously cryptic gene cluster coding for the biosynthesis of the lanthipeptide nostolanthin (*nlt*), which is rapidly induced upon physical contact between *Nostoc punctiforme* and the liverwort *Blasia pusilla*. Despite its robust transcriptional activation, nostolanthin remained undetectable in its native producer by conventional metabolomic analyses. Heterologous reconstitution of the biosynthesis showed that the lanthipeptide synthetase NltM produces a bicyclic class II lanthipeptide containing a non-cyclized dehydroamino acid together with a free cysteine residue. We show that this lanthipeptide undergoes covalent oligomerization into high-molecular-weight assemblies, a process favored under native cyanobacterial expression conditions. An antibody raised against the oligomeric peptide enabled the detection of secreted nostolanthin in cyanobacteria and revealed that it accumulates predominantly in an oligomeric form. Comparative genomics showed that nostolanthin belongs to a larger family of Nif11-type-lanthipeptide biosynthetic gene clusters, consistently associated with homologous two-component regulatory systems. Importantly, oligomeric, but not monomeric nostolanthin, accelerated the establishment of physical contact between *Nostoc* and *B. pusilla*. Together, these findings reveal oligomerization by covalent bond formation as a previously unrecognized mode of lanthipeptide maturation and identify nostolanthin as a host-responsive peptide that regulates the transition from a free-living to a symbiotic lifestyle.

## Introduction

*Nostoc* is a key nitrogen-fixing cyanobacterial symbiont in terrestrial ecosystems, forming associations with lichens, bryophytes, and some higher plants. Through these partnerships, it enhances host growth in nutrient-poor soils and contributes significantly to overall ecosystem productivity.^1–3^ This filamentous, heterocyst-forming genus includes both free-living and symbiotic strains with a broad host range. It occurs in extracellular and intracellular associations, often involving the development of specialized host structures that accommodate the cyanobacteria.^2^

Symbiotic *Nostoc* strains have a polyphyletic origin, with most strains clustering within host-generalist lineages that associate with a wide range of partners, including bryophytes, lichens, cycads, and *Azolla*. ^4,5^ This ecological versatility points to diverse strategies for establishing host-specific interactions. Phylogenomic studies of *Nostoc* reveal enrichment of particular functional gene categories in symbiont clades, notably biosynthetic gene clusters (BGCs) involved in a specialized metabolism. ^4,6^ A recent meta-analysis of BGC abundance and diversity identified *Nostoc* as the most prolific genus within the phylum Cyanobacteria. ^7,8^ Indeed, numerous structurally unique natural products have already been identified in this genus. ^9^ Most characterized metabolites are derived from non-ribosomal peptide synthetase (NRPS) or polyketide synthase (PKS) pathways, with additional compounds belonging to ribosomally synthesized and posttranslationally modified peptide (RiPP) or terpene classes. ^9–11^ Despite these advances, the majority of BGCs remain uncharacterized, highlighting a vast untapped biosynthetic potential and significant gaps in our understanding of their function. ^7^

In contrast to extensive genomic and evolutionary work on *Nostoc*, functional studies on the involvement of secondary metabolites particularly in symbiosis remain limited. It is known that *Nostoc* in response to host factors forms hormogonia – motile non-growing filaments that are essential for penetrating the hosts ^12,13^. These hormogonia-inducing factors (HIFs) are counterbalanced by hormogonia-repressing factors (HRFs), which can influence the extent of chemoattraction from the cyanobacterial side and also stimulate the reversion of hormogonia inside plants. ^2^ In *Nostoc punctiforme* PCC 73102, the NRPS-derived peptide nostopeptolide has been identified as a key repressor of hormogonia formation. ^14^ Emerging evidence further shows that specific BGCs are transcriptionally activated during host interactions – for example, a polyketide BGC and its associated FG-GAP protein are upregulated in the epiphytic interaction with the feather moss *Pleurozium schreberii*. ^15^ A functional role of small molecules in symbiotic interactions of *Nostoc* is also supported by metabolomic studies; for example, MALDI imaging of *Nostoc* inside the host plant *Gunnera manicata* revealed both upregulation and downregulation of mostly cryptic small molecules. ^14^ In another study, a comparison of *Nostoc* isolates from *Blasia pusilla* and the surrounding soil provided evidence of a possible discriminating role of small molecules in symbiotic recruitment. ^16^

Against this backdrop, however, the contribution of specific classes of specialized metabolites remains unevenly explored. In particular, RiPPs – despite their biosynthetic diversity and prevalence in cyanobacteria – have received comparatively little attention in the context of symbiotic interactions. Although several cyanobacterial RiPP families – such as microviridins, cyanobactins, nostatin, prochlorosins, and synechococsins – have been described, their functional roles remain largely unclear. ^10,17–20^ Some may contribute to defend against competitors or grazers, as suggested for synechococsins and the highly cytotoxic nostatin. ^10,17^ Several additional cyanobacterial RiPP pathways have been reconstructed in *Escherichia coli*, revealing unique peptide modifications, ^21–23^ but the corresponding peptides have so far eluded detection in the native producers.

Here, we characterize a cryptic lanthipeptide BGC that is rapidly and strongly upregulated during physical interaction with the bryophyte host *B. pusilla*. Although the product, nostolanthin, was successfully reconstituted in *E. coli*, it was not detectable in cyanobacterial hosts, even upon strong overexpression of its biosynthetic genes. Our data suggest that the cyanobacterial background promotes covalent oligomerization of the peptide, thereby rendering it undetectable by standard analytical techniques. Functionally, nostolanthin overexpression promoted earlier physical contact between *Nostoc* and *B. pusilla*, indicating a role in early adaptation to the transition from a free-living to a symbiotic lifestyle.

## Results

### Physical interaction of *Nostoc* and *Blasia* induces cryptic lanthipeptide gene expression but no detectable lanthipeptide production

To evaluate the potential role of specialized metabolites in the interaction between *N. punctiforme* and the model host *B. pusilla*, transcriptional profiling was employed to assess the expression of 14 characterized and cryptic NRPS, PKS, and RiPP BGCs (see previous BGC nomenclature ^24^). *B. pusilla* was pre-cultivated under nitrogen-deprived conditions, after which *N. punctiforme* was either exposed to the resulting exudate (Fig. 1a) or co-cultivated with the starved liverwort (Fig. 1b, dataset S1). Exposure to *B. pusilla* exudate had only a modest effect on expression of representative biosynthetic genes, which were selected based on our previous RNA sequencing study^18^ (Table S3C): among the clusters analyzed, only the two cryptic RiPP BGCs, *ripp3* and *ripp4*, exhibited transient upregulation after 1 h, which subsided by 24 h (Fig. 1c). In contrast, the physical interaction between *B. pusilla* and *N. punctiforme* triggered a markedly stronger response, with *ripp3* and *ripp4* showing a pronounced transcriptional upregulation that likewise declined after 24 h (Fig. 1c). At this later time point, several additional BGCs only displayed moderate induction, including the *pks5* cluster, which has previously been reported to respond to interactions with feather mosses (Fig. 1c). ^15^ Based on these observations, the *ripp4* BGC was selected for further analysis. To validate and visualize its transcriptional response to *B. pusilla*, we employed a recently developed *ripp4* cyan fluorescent protein (CFP)-based reporter strain of *N. punctiforme* which harbors a replicative plasmid carrying the BGC’s 5’ untranslated region (UTR) fused to CFP, allowing transcriptional activity to be monitored via CFP signal. ^18^ This analysis confirmed a strong induction of the *N. punctiforme ripp4* reporter strain upon physical interaction of the partners, detectable as early as 1.5 h (Fig. 1d). Notably, the reporter signal remained robust even after 24 h. Overlay of CFP fluorescence with cyanobacterial autofluorescence further revealed pronounced phenotypic heterogeneity, with approximately half of the filaments exhibiting strong *ripp4*-associated fluorescence (Fig. 1d).

**Fig. 1.**
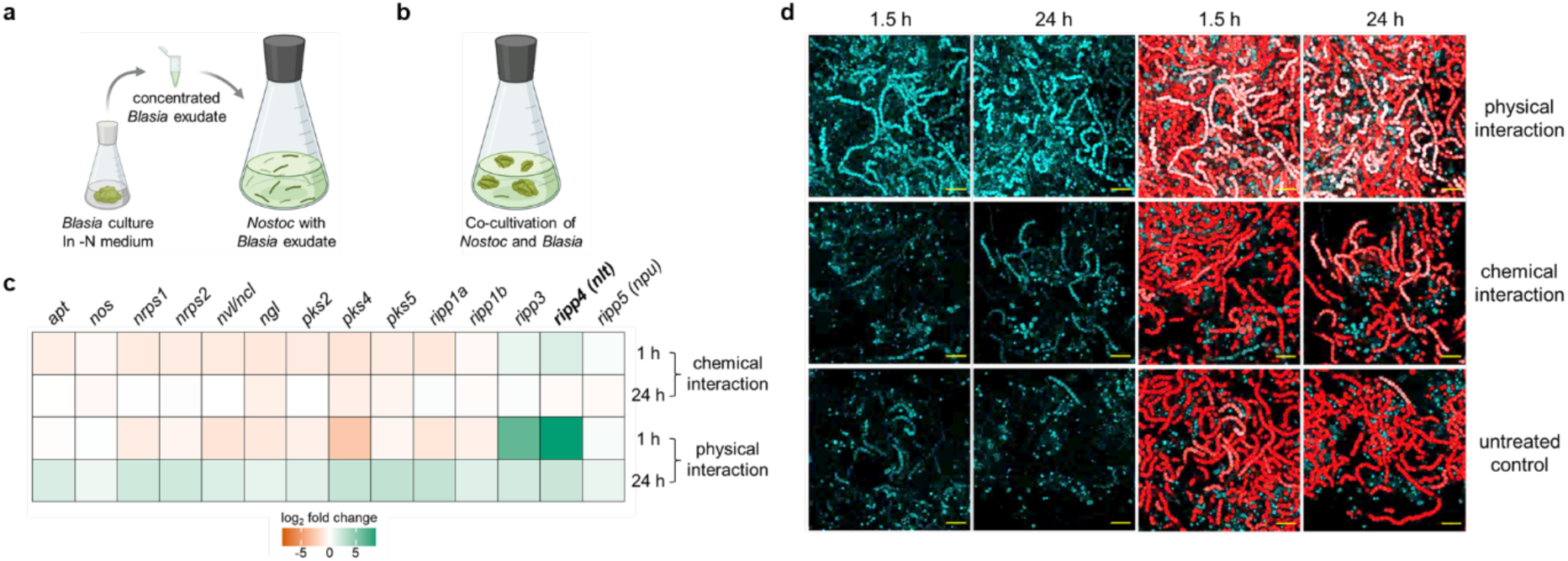
Interaction of *N. punctiforme* with the host plant *Blasia pusilla*. **a** Schematic representation of chemical interaction induced by addition of concentrated *B. pusilla* exudate to a nitrogen-starved *N. punctiforme* culture. **b** Schematic representation of physical interaction induced by addition of *B. pusilla* thalli to a nitrogen-starved *N. punctiforme* culture. **c** RT-qPCR analysis of RNA collected 1 h and 24 h after initiation of the interaction to determine transcription levels of representative genes associated with secondary metabolite biosynthetic gene clusters (BGCs) of *N. punctiforme* (Table S3C). Values represent average fold changes of biological triplicates relative to an untreated control culture (Dataset S1). **d** Confocal fluorescence micrographs of the *N. punctiforme ripp4* reporter strain harboring a replicative plasmid carrying a 614 bp 5’ UTR of the *ripp4* BGC fused to a cyan fluorescent protein (CFP). ^18^ Co-cultivation with *B. pusilla* or *B. pusilla*-conditioned exudate for up to 24 h, relative to an untreated control. Left: CFP channel signal at 1.5 h and 24 h of cultivation (green). Right: merged CFP and Chl*α* autofluorescence channel signals at the corresponding time points (red). Scale bar: 20 µm.

Both *ripp3* and *ripp4* BGCs encode class II lanthipeptides and include a LanM-type lanthipeptide synthetase, which typically catalyzes both the dehydration of serine and threonine residues and the subsequent intramolecular thioether bond formation, yielding lanthionine or methyllanthionine crosslinks between cysteines and the corresponding dehydroamino acids (Fig. 2a). Based on this annotation, the putative *ripp4* product was designated as nostolanthin (*nlt*) BGC. Using the inferred sequence of the unmodified core peptide as a query, we searched for nostolanthin and potential derivatives using both targeted and untargeted metabolomics approaches; however, no candidate compounds could be detected in either mono-or co-cultures (Supplementary Fig. 1).

**Fig. 2.**
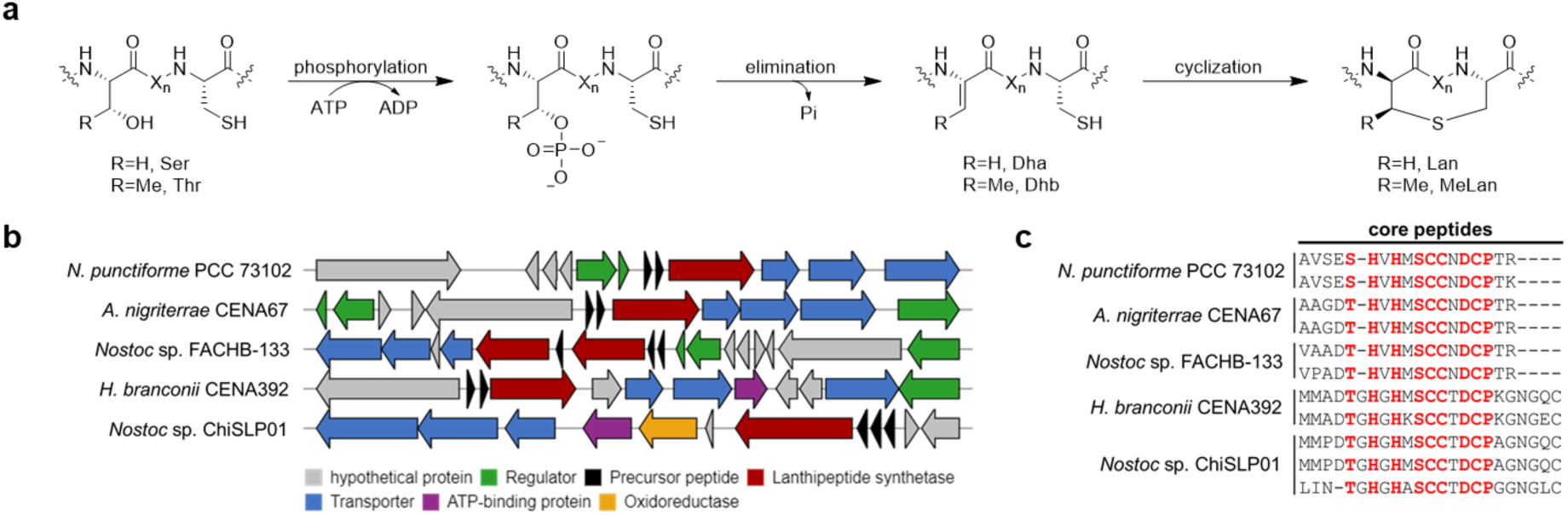
Nostolanthin is a class II lanthipeptide encoded by a BGC distributed across filamentous, nitrogen-fixing cyanobacteria. **a** Biosynthesis of class II lanthipeptides, Dha: Dehydroalanine, Dhb: Dehydrobutyrine, (Me-)Lan: (Methyl-)Lanthionine. **b** The nostolanthin (*nlt*) BGC (formerly designated *ripp4*) from *N. punctiforme* PCC 73102 and representative, closely related BGCs identified in other filamentous, nitrogen-fixing cyanobacteria based on a BLASTP search of the precursor peptides and the lanthipeptide synthetase. **c** Alignment of the corresponding core peptides show a conserved motif, highlighted in red.

### Phylogenetic Distribution and Conserved Architecture of the *nlt* Biosynthetic Gene Cluster

A BLASTP search identified close homologs of nostolanthin-like precursor peptides NltA associated with an NltM-like lanthipeptide synthetase in 28 cyanobacterial strains (as of August 2026). These homologs are exclusively encoded within highly similar BGCs from filamentous, nitrogen-fixing cyanobacteria belonging to the order *Nostocales* (Fig. 2b). The identified BGCs display a conserved genetic architecture, comprising one or two class II lanthipeptide synthetase genes, a pair of precursor peptide genes, and three transporter genes, closely resembling the organization of the *nlt* BGC.

Each nostolanthin-like BGC encodes two precursor peptides, NltA1 and NltA2, each consisting of a Nif11-like leader peptide followed by a core peptide containing three to four serines or threonines and three cysteine residues (Fig. 2c). Sequence alignment revealed a highly conserved core peptide motif, (S/T)HxHxSCCxDCP, suggesting that this region is critical for the structure and/or function of this distinct family of lanthipeptides (Fig. 2c). Notably, the precursor peptides from *Halotia branconii* CENA392 and *Nostoc* sp. ChiSLP01 possess an extended C terminus containing an additional cysteine residue, extending the core peptide length from 19 to 24 amino acids (Fig. 2c). All leader peptides contain a conserved double-glycine motif, indicative of a leader peptide cleavage site by a C39 peptidase. Consistent with this prediction, all identified BGCs encode three putative transporter proteins (NltT1–T3), with NltT2 harboring an N-terminal C39 peptidase domain. In addition to the core biosynthetic genes, the *nlt* BGC encodes a two-component regulatory system comprising a multidomain sensor kinase NltR1 with a histidine kinase-like ATPase domain and a cognate response regulator NltR2 (Fig. 2b and Table S4). This regulatory module may sense host-derived signals and activate expression of the *nlt* BGC. To investigate whether this regulatory module is conserved among symbiotic *Nostoc* strains, we performed an additional BLASTP and CluSEEK ^25^ analysis of the response regulator, which contains a PAS-type sensory domain. ^26^ This analysis revealed that the *nlt* BGC belongs to a larger and widespread family of Nif11-type RiPP BGCs present in numerous *Nostocales* strains (57 as of August 2026), all associated with a homologous two-component regulatory system and a LanM-type lanthipeptide synthetase. Although these BGCs share a highly conserved genetic organization, the encoded core peptides differ substantially in their primary sequence (Supplementary Fig. 2). Interestingly, some of these nostolanthin-like BGCs also encode internalins, a family of leucine-rich repeat proteins known to mediate contact-dependent host interactions in *Listeria* ^27^ (Supplementary Fig. 2). These findings suggest that nostolanthine-like RiPPs may be part of a larger molecular machinery involved in contact-dependent interactions with the host, in this case *B. pusilla*.

### NltM catalyzes the formation of a two-ring lanthipeptide with a remaining dehydroamino acid-cysteine pair in *E. coli*

To investigate the catalytic activity of NltM, the N-terminally His-tagged precursor peptide NltA1 was co-expressed with the synthetase NltM in *E. coli* using a Duet vector system (Fig. 3a). Following purification of the lysate by immobilized metal ion affinity chromatography (IMAC) and *in vitro* leader peptide cleavage with LahT150, a C39 peptidase homologous to the N-terminal peptidase domain of NltT2, LC-MS analysis revealed a threefold dehydrated core peptide with a mass of 1920.75 Da (3x H_2_O = –54.03 Da; Fig. 3b). Lanthionine ring formation was assessed by iodoacetamide (IAA) alkylation of free cysteine residues. Detection of a single IAA adduct (+57.02 Da; Fig. 3b) repeatedly indicated that one cysteine remained uncyclized, whereas the other two were engaged in thioether cross-links, consistent with the formation of two lanthionine rings. The presence of a threefold dehydration alongside two lanthionine rings necessarily implies the existence of one unreacted dehydroamino acid residue within the core peptide. This is further supported by the detection of a highly abundant species at 2227.83 Da, consistent with a glutathione (GSH) adduct (+307.08 Da, Supplementary Fig. 3) to the dehydroamino acid, an artefact that has been previously observed during heterologous expression of lanthipeptides in *E. coli*. ^28^ To determine the amino acid residues modified by NltM, LC-MS/MS analysis of the core peptide mass was performed. Analysis of the fragment ions revealed that Ser5, Ser10 and Thr17 underwent dehydration to Dha5, Dha10 and Dhb17, respectively (Fig. 3c). Furthermore, the fragmentation pattern suggests that the two lanthionine rings formed are overlapping, leaving Dhb17 as the unreacted dehydro amino acid.

**Fig. 3.**
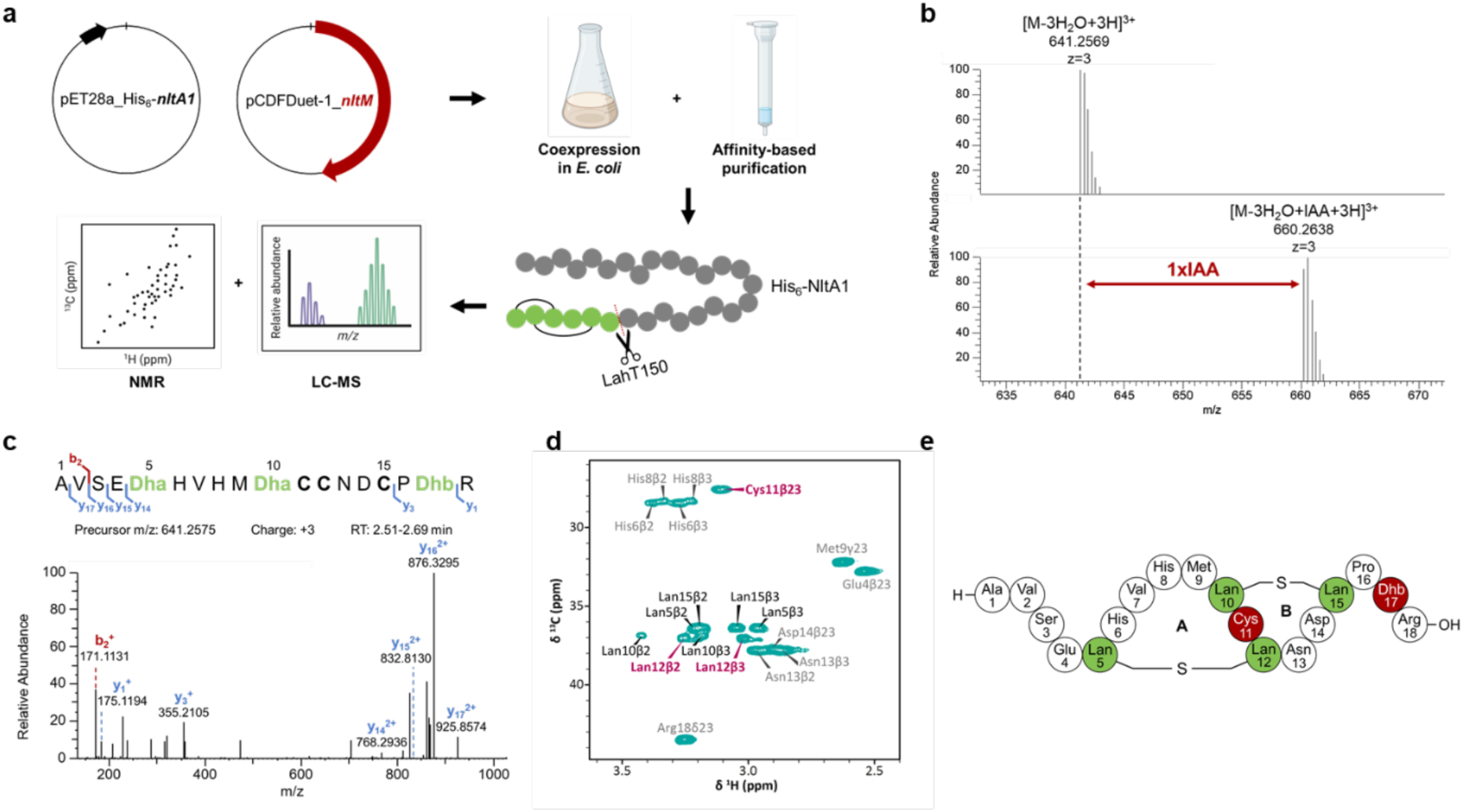
Characterization of NltM-modified His₆-NltA1 produced in *E. coli*. **a** Workflow for *in vivo* reconstitution of the *nlt* BGC. His₆-NltA1 and NltM were co-expressed in *E. coli* LOBSTR cells, the peptide was purified by affinity chromatography, the leader peptide was cleaved using the C39 peptidase LahT150, and the modified core peptide was analyzed by LC-MS and NMR spectroscopy. **b** LC-MS spectra of LahT150-cleaved His₆-NltA1 before (top) and after (bottom) iodoacetamide (IAA) treatment. Top: [M-3H₂O+3H]³⁺, calculated: 641.2569, observed: 641.2569. Bottom: [M-3H₂O+IAA+3H]³⁺, calculated: 660.2635, observed: 660.2638. **c** LC-MS/MS spectrum of LahT150-cleaved His₆-NltA1 at *m/z* 641.2575, with assigned b-and y-ions. **d** Diagnostic Cβ region of the 2D ^1^H-^13^C HSQC spectrum of nostolanthin-T17A in water at 308 K. The pairs of methylene cross peaks of lanthionine residues are labeled in black, whereas those of residues Cys11 and Lan12 are highlighted in red. The Cβ chemical shift is diagnostic for the redox state of the neighboring sulfur atom (approx. 27 ppm for reduced thiol of Cys11 versus 37 ppm for the oxidized thioether of Lan12). **e** Chemical structure of the lanthipeptide nostolanthin A (1920.75 Da). The residues involved in the two lanthionine rings are highlighted in green, the uncyclized Cys11 and the unreacted Dhb17 are highlighted in red.

To further investigate the precursor peptide processing and the overlapping ring topology, site-directed mutagenesis of NltA1 was performed to generate the four variants NltA1-S5A, NltA1-C11A, NltA1-C12A, and NltA1-T17A. All mutant precursor peptides were co-expressed with NltM, purified as described above, subjected to *in vitro* leader peptide cleavage, and analyzed by IAA alkylation and LC-MS/MS. The nostolanthin-S5A variant underwent only two dehydrations, confirming Ser5 as a substrate for NltM-catalyzed dehydration. Consistent with the loss of one lanthionine linkage, two IAA adducts were detected, indicating the presence of two free cysteine residues. LC-MS/MS analysis further supported the formation of a single lanthionine ring, with the fragmentation pattern suggesting a thioether bridge between Ser10 and Cys15 (Supplementary Fig. 4-5). However, the topology of the second ring could not be unambiguously resolved from these data alone. In contrast, both nostolanthin-C11A and nostolanthin-C12A variants exhibited three dehydrations and no IAA adducts, indicating full involvement of the cysteines in the formation of two lanthionine rings (Supplementary Fig. 6-9). These results demonstrate that either Cys11 or Cys12 can substitute for the other in forming the second lanthionine linkage with Ser5. The nostolanthin-T17A variant exhibited only two dehydrations and a single IAA adduct, confirming Thr17 as the third dehydration site while indicating one free cysteine, as observed for the wild-type peptide (Supplementary Fig. 10-11). Notably, unlike the wild type, nostolanthin-T17A did not form a GSH adduct, further supporting the conclusion that Dhb17 remains as an uncyclized dehydroamino acid and serves as the site of GSH addition in the heterologous producer, rather than participating in lanthionine ring formation.

To definitively determine whether Cys11 or Cys12 forms the second lanthionine, the modified peptide was subsequently subjected to nuclear magnetic resonance (NMR) spectroscopy. For this structural analysis, we used the maturated nostolanthin-T17A variant to avoid glutathionylation. The peptide was dissolved in water and sequentially assigned using 2D NMR spectra (see methods section, Supplementary Fig. 12-14, and Table S5). The resonance assignment unambiguously confirmed the peptide constitution comprising two lanthionine rings: one thioether bridge is established between residues Ser10 and Cys15, as suggested from MS data, and the second lanthionine ring involves residues Ser5 and Cys12. The free thiol of residue Cys11 gave rise to a diagnostic ^13^Cβ chemical shift of 27.6 ppm, whereas all methylene carbon atoms of the thioether linkages resonated at characteristic 36-37 ppm (Fig. 3d). Together with sequential Hα/β_i_-HN_i+1_ contacts in the NOESY spectrum, in particular between Lan12 and Asn13, the NMR and MS data excluded other ring topologies (Fig. 3e). Purified nostolanthin tested against Gram-positive bacteria, Gram-negative bacteria, and cyanobacteria, did not show inhibitory activity (Supplementary Fig. 15).

### Native overexpression of the NltA1 precursor peptide uncovers an unexpected oligomeric state

To verify NltM-dependent modifications in the native cyanobacterial environment, NltA1 was overexpressed in *N. punctiforme* as an N-terminally His-tagged precursor peptide to enable its purification and characterization. For this purpose, *nltA1* was cloned under the control of a strong constitutive promoter previously used for heterologous gene expression in *N. punctiforme* on a replicative plasmid. ^29^ To prevent leader peptide cleavage during maturation, the conserved Gly-Gly motif was replaced with Ile-Ile, thereby allowing purification of the intact modified precursor peptide by affinity chromatography (Fig. 4a). The resulting construct was introduced into *N. punctiforme* by transformation. Unexpectedly, immunoblot analysis with a monoclonal anti-polyhistidine antibody following affinity purification did not detect the expected 11.4 kDa precursor peptide. Instead, a prominent band was observed at approximately 250 kDa, suggesting that NltA1 forms a high-molecular-weight oligomer or aggregate (Fig. 4b). This species remained stable under denaturing and reducing conditions (8 M urea), suggesting possible covalent cross-links (Fig. 4b). The high-molecular-weight species did not yield a distinct MS/MS pattern and therefore could not be further analyzed with regard to its modification(s).

**Fig. 4.**
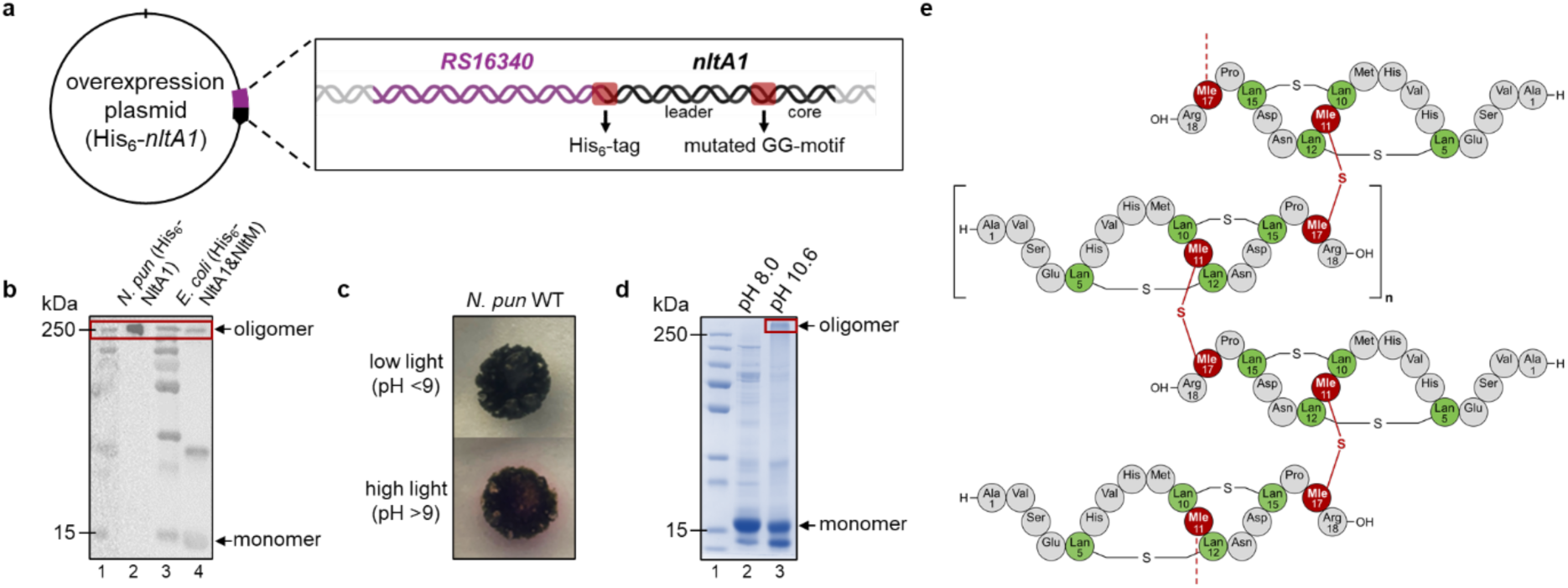
Oligomerization of the NltA precursor peptide in *N. punctiforme* and *E. coli*. **a** Schematic of the overexpression plasmid (pRL1049-nltA1), harboring the strong constitutive promoter *NPUN_RS16340* and the precursor peptide gene *nltA1*, modified to encode an N-terminal His-tag and a mutated GG-motif to enable purification of the modified, full-length precursor peptide for subsequent analysis of its modifications (used in *N. punctiforme*). **b** Western blot analysis of the overexpressed peptide purified from the *N. punctiforme* mutant harboring the vector pRL1049-nltA1 by affinity chromatography under denaturing conditions using 8 M urea (lane 2) and the peptide co-expressed with NltM and purified from *E. coli* by affinity chromatography under native conditions (lane 4). Lane 1 and 3 show the protein ladder. **c** *N. punctiforme* cultures grown on BG11 agar plates supplemented with phenolphthalein (0.01 mg/mL) under low-light (12 µE; top) and high-light (75 µE; bottom) conditions. **d** SDS-PAGE of *E. coli*-derived, affinity-purified precursor peptide incubated for 48 h at room temperature in buffer at pH 8 (peptide lysis buffer, lane 2) versus pH 10.6 (carbonate-bicarbonate buffer, lane 3). The high-molecular-weight species was enriched by analytical size-exclusion chromatography (V_e_: 8.86 mL) (Supplementary Fig. 16). **e** Structural model of the oligomeric nostolanthin A. The lanthipeptide is proposed to form intermolecular Methyllanthionine (Mle) bridges involving Cys11 and Dhb17, leading to covalent oligomerization of the mature peptide.

Re-examination of the NltA1/NltM co-expression system in *E. coli* likewise revealed, in addition to monomeric precursor peptide with a molecular weight of 11.9 kDa, a high-molecular-weight band of approximately 250 kDa (Fig. 4b). Although these expression systems are not directly comparable, the preferential formation of the high-molecular-weight species under the cyanobacterial expression conditions suggested that host-specific factors may promote oligomerization. We hypothesized that the more alkaline extracellular environment of cyanobacteria, which is the result of photosynthesis and CO_2_ fixation, ^30^ could enhance oligomerization. To test this, wild-type *N. punctiforme* cultures were grown on agar plates supplemented with phenolphthalein under either low light conditions (12 µmol photons m^-2^s^-1^) to minimize potential effects of photosynthesis or alternatively high light conditions (75 µmol photons m^-2^s^-1^) to promote optimal photosynthesis. Under high-light conditions, the extracellular region surrounding the culture turned pink, indicating a pH > 9 under conditions stimulating photosynthesis (Fig. 4c). To determine whether alkaline conditions promote precursor peptide oligomerization, the *E. coli*-derived affinity-purified NltA1 precursor was incubated for 2 days at pH 8 and pH 10.6, respectively. SDS-PAGE analysis revealed a decrease in monomeric precursor peptide at pH 10.6 accompanied by accumulation of the high-molecular-weight species, demonstrating that an elevated pH indeed promotes oligomer formation (Fig. 4d). The relatively modest extent of conversion likely reflects the presence of glutathionylated precursor peptide generated during *E. coli* expression (Supplementary Fig. 3), which is unavailable for pH-dependent cross-linking. The high-molecular-weight species was subsequently enriched by size-exclusion chromatography (SEC). Comparison with SEC standards supported a molecular weight > 200 kDa (Supplementary Fig. 16). The purified high-molecular-weight form of nostolanthin, which retained the uncleaved Nif11 leader peptide, was subsequently used as antigen to generate the polyclonal antibody anti-NltA1.

### Oligomeric nostolanthin influences cyanobacterial motility and symbiotic interactions

To investigate the functional role of nostolanthin in symbiotic interactions, while enabling tag-free purification of the peptide under native cyanobacterial expression conditions, we adopted a gain-of-function strategy by introducing the complete nostolanthin BGC into the symbiotic strain *Nostoc* sp. KVJ2, which was originally isolated from *B. pusilla* and lacks an endogenous *nlt* BGC. ^31^

For this purpose, the *nlt* BGC was placed under the control of a constitutive promoter on a replicative plasmid as described previously (^29^, Fig. 5a) and introduced into *Nostoc* sp. KVJ2 by transformation. RT-qPCR confirmed transcription of the introduced BGC in KVJ2-*nlt* (Supplementary Fig. 17); however, LC-MS analysis repeatedly failed to detect the monomeric peptide product. Because the un-tagged, putative oligomeric product could not be detected by mass spectrometry or standard proteomic approaches, the newly generated polyclonal antibody anti-NltA1 was employed for its detection. Initial tests of the capabilities of the antibody showed that it recognized both, monomer and high-molecular-weight forms of the nostolanthin precursor in *E. coli* (Supplementary Fig. 18). As immunoblotting of *Nostoc* cell extracts and supernatant extracts on SDS-PAGE gels produced diffuse signals, comparative dot-blot analysis was performed on pellet and supernatant fractions from wild-type and KVJ2-*nlt* strains. A weak signal was detected in the pellet fraction of both strains, potentially due to cross-reactivity with endogenous Nif11-like peptides, whereas a strong signal was observed exclusively in the supernatant of the KVJ2-*nlt* strain (Fig. 5b). No corresponding signal was detected in the supernatant of the wild type. Further fractionation of the mutant supernatant by SEC, followed by dot-blot analysis of individual fractions, revealed a signal confined to a single high-molecular-weight fraction (V_e_ = 8.34 mL), corresponding to the exclusion volume of the column (upper molecular weight limit, approx. 600 kDa). This supports the hypothesis that the secreted nostolanthin assembles into a covalent peptide oligomer in *Nostoc* sp. KVJ2 (Fig. 5c). However, the amount of the oligomeric nostolanthin species was too low for further analytical characterization.

**Fig. 5.**
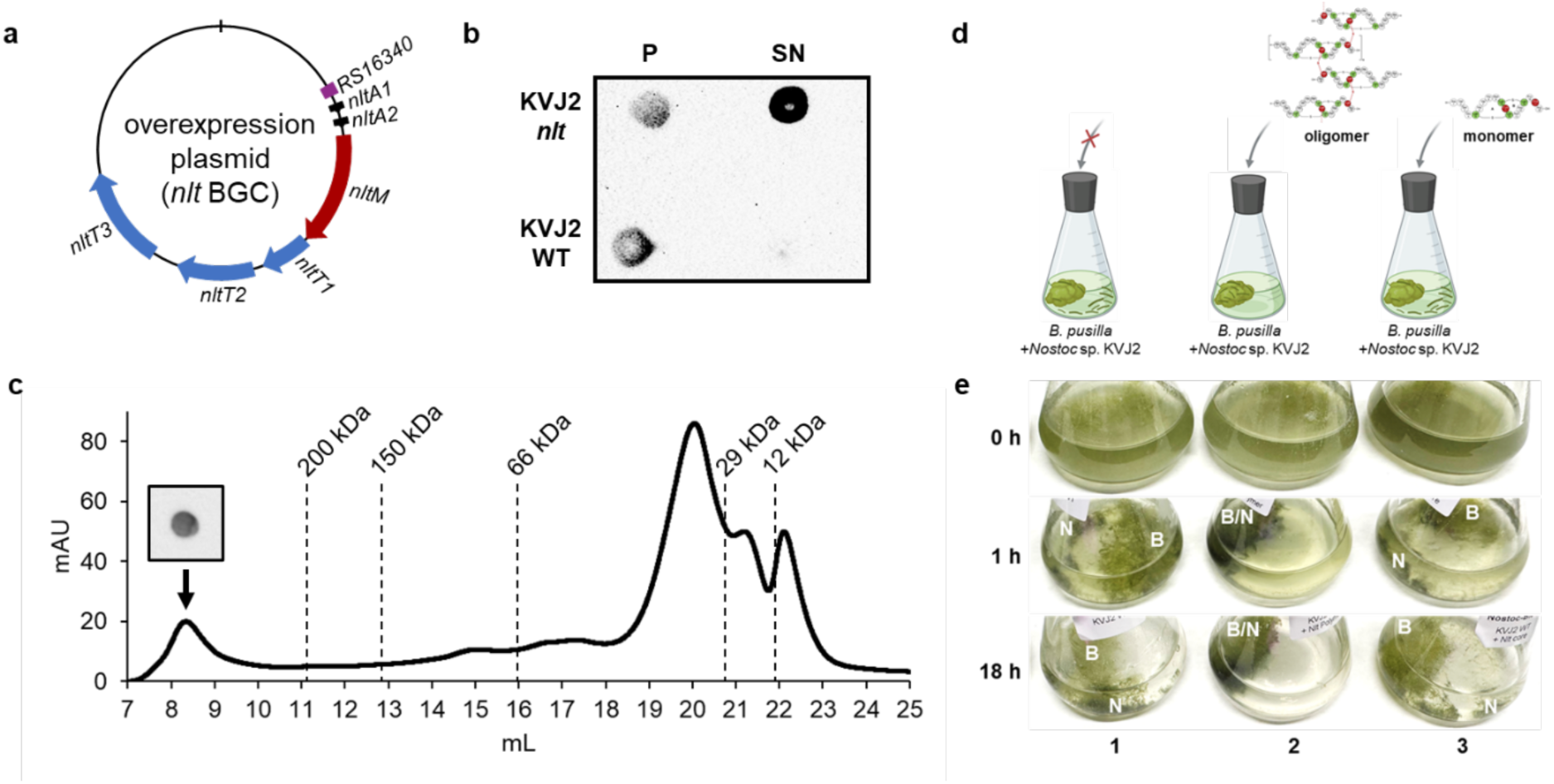
Characterization of the *Nostoc* sp. KVJ2 mutant strain harboring the *nlt* BGC. **a** Schematic of the overexpression plasmid (pRL1049-nlt_BGC), harboring the strong constitutive promoter *NPUN_RS16340* and the *nlt* BGC. **b** Dot-blot comparison of pellet (P) and supernatant (SN) fractions from *Nostoc* sp. KVJ2 WT and KVJ2-*nlt* cultures employing the polyclonal antibody anti-NltA1 targeting the monomer and high-molecular-weight forms of the nostolanthin precursor purified from *E. coli*. **c** SEC profile of the *Nostoc* sp. KVJ2-*nlt* supernatant, obtained using a Superdex 200 Increase 10/300 GL column (bed volume: 24 mL). The peak at RT 8.34 mL corresponds to the fraction giving a signal in the dot-blot using the anti-NltA1 antibody. This elution volume coincides with the exclusion limit of the column (approx. 600 kDa). **d** Schematic representation of the *Nostoc-B. pusilla* symbiosis assay. *Nostoc* sp. KVJ2 wild-type was added to liquid cultures of nitrogen-starved *B. pusilla*. The nostolanthin oligomer purified from the KVJ2-*nlt* supernatant and the nostolanthin monomer purified from *E. coli* were added to separate flasks; a control without the addition of a peptide was included. **e** Photographs of *Nostoc*-*B. pusilla* symbiosis assays. The interaction between nitrogen-starved *B. pusilla* and *Nostoc* sp. KVJ2 was assessed under three conditions: control (1), addition of the nostolanthin oligomer (2), and addition of the nostolanthin monomer (3). Images were taken at 0 h, 1 h, and 18 h after addition of *Nostoc* sp. KVJ2 and the respective peptides. Addition of the nostolanthin oligomer resulted in rapid association between *B. pusilla* (B) and *Nostoc* sp. KVJ2 (N). In contrast, in the absence of the nostolanthin oligomer, *Nostoc* biomass remained partially dispersed in the medium with only limited accumulation around the liverwort.

To investigate the role of the *nlt* BGC in the symbiotic interaction between *B. pusilla* and *Nostoc*, wild-type *Nostoc* sp. KVJ2 was co-cultivated with nitrogen-starved *B. pusilla* cultures under three conditions: addition of SEC-purified nostolanthin oligomer isolated from KVJ2-*nlt* culture supernatant, addition of monomeric nostolanthin A (wild-type core peptide) purified after heterologous expression in *E. coli*, and an untreated control (Fig. 5d). Within 1 h, cultures supplemented with nostolanthin oligomer showed markedly accelerated physical association between the two partners, with cyanobacterial filaments and liverwort tissue in very close contact (Fig. 5e). This effect became even more pronounced after 18 h, at which point the culture medium had cleared almost entirely of dispersed cyanobacterial biomass. In contrast, both the untreated control and the monomer-treated cultures showed only partial association at this time point, with substantial cyanobacterial biomass remaining dispersed in the medium (Fig. 5e). Together, these results indicate that the nostolanthin oligomer, but not its monomeric form, accelerates the establishment of physical contact between *Nostoc* and *B. pusilla*, supporting a role for this lanthipeptide in the early stages of symbiotic interaction.

## Discussion

This study provides new insights into the structure and function of a previously enigmatic family of lanthipeptides in *Nostoc* termed nostolanthines. We demonstrate that production of nostolanthin is strongly induced upon physical host contact and facilitates the establishment of physical associations between *N. punctiforme* and *B. pusilla* during the initial phase of symbiosis. In addition, we uncover a previously unrecognized consequence of lanthipeptide biosynthesis: the covalent oligomerization of the mature peptide. This process is strongly promoted under native cyanobacterial expression conditions and has likely obscured the detection and characterization of an initially expected monomeric nostolanthin to date.

The nostolanthin BGC is largely restricted to filamentous, nitrogen-fixing cyanobacteria, many of which establish symbiotic associations with plants. However, its distribution is sporadic, raising questions about the specificity of its function and its broader role in symbiosis. Notably, we found that the combination of homologous sensor-response regulator pairs and Nif11-type lanthipeptide BGCs is widespread among symbiotic *Nostoc* strains. One possibility is that the different nostolanthin subfamilies confer specificity toward distinct host species. This hypothesis could be tested by comparing the symbiotic performance of strains producing different nostolanthin variants in association with *B. pusilla* and other compatible hosts. Alternatively, the primary sequence of the lanthipeptides may be less important for host interaction rather than the higher-order structure generated through covalent oligomerization. Under this scenario, sequence diversification of the core peptides would primarily reflect the remarkable evolutionary plasticity of cyanobacterial lanthipeptides ^32^ rather than functional specialization. This structural hypothesis is further supported by the observation that several nostolanthin-like BGCs also encode putative internalins. ^27^ In *Listeria*, these leucine-rich repeat proteins adopt a characteristic concave horseshoe-shaped structure that mediates binding to host cadherins, thereby facilitating intimate cell-cell interactions. ^27^ By analogy, the co-occurrence of internalins and nostolanthin-like lanthipeptides raises the possibility that these molecules cooperate in mediating contact-dependent interactions between *Nostoc* and its plant hosts. It can be further speculated that immobilization of oligomeric nostolanthin and the use of a nostolanthin-specific antibody may enable the identification of interacting proteins in *Blasia* and *Nostoc*. Such strategies, including pull-down assays followed by protein identification, may help reveal potential binding partners and provide mechanistic insights into the function and mode of action of nostolanthin.

Nostolanthins are not the only cyanobacterial lanthipeptides that have thus far evaded detection in their native hosts. Notably, cyanobacterial lanthipeptides frequently contain dehydroamino acids that are not engaged in intramolecular thioether ring formation. This is exemplified by lanthipeptides NpnA3 and NpnA6 that fully lack cysteine residues and instead comprise several unlinked dehydroamino acids ^33^, making them particularly susceptible to intermolecular reactions with cysteine-containing peptides or other nucleophilic moieties of cellular components. Although the corresponding RiPP BGCs are actively transcribed ^18^, their products have so far escaped analytical detection in *N. punctiforme*. We therefore propose that homo-or heterooligomeric RiPP assemblies may be far more widespread in cyanobacteria than previously anticipated. The factors driving this apparent propensity for oligomerization remain unclear. It is conceivable that the photosynthesis-induced alkaline extracellular environment of cyanobacteria ^30^ favors spontaneous intermolecular Michael additions, although enzymatic crosslinking cannot currently be fully excluded. Because pH values above 8 increase the abundance of thiolate species by shifting the thiol-thiolate equilibrium ^34^, the resulting higher nucleophilicity may promote oligomerization through Michael addition reactions. ^35^ In addition, the comparatively lower ratio of reduced glutathione (GSH) to oxidized glutathione (GSSG) in cyanobacteria ^36^ relative to *E. coli* ^37^ may differentially influence the availability and reactivity of the free cysteine residue and the dehydrobutyrine (Dhb) moiety of nostolanthin. These factors may explain why the peptide was readily detected as a monomer and glutathione adduct in *E. coli* but not in its native cyanobacterial producer. While this extensive oligomerization initially posed a major analytical challenge that hindered the functional characterization of nostolanthin, it ultimately proved advantageous. By increasing the pH of *E. coli* fractions, we were able to promote oligomer formation *in vitro*, isolate high-molecular-weight oligomers by size-exclusion chromatography, and use the purified material as an antigen for antibody production. The resulting antibody enabled the detection of mature, leader-peptide-free high-molecular-weight RiPP oligomers in their native context without the need for affinity tags or mass spectrometric analyses. By contrast, antibody generation against monomeric lanthipeptides typically requires conjugation of the low-molecular-weight peptides to carrier proteins. For example, a recently reported paenibacillin antibody was generated using the peptide conjugated to keyhole limpet hemocyanin (KLH). ^38^ In contrast, purified oligomeric nostolanthin itself elicited a robust immune response in rabbits, enabling the generation of antibodies that reliably recognize the native peptide assemblies.

We were ultimately able to demonstrate that the addition of oligomeric nostolanthin directly promotes the physical interaction between *Blasia* and *Nostoc*, providing direct functional evidence that these peptides facilitate contact-dependent interactions. Elucidating the underlying molecular mechanism will require future studies of protein-protein interactions and structural analyses, both of which depend on the large-scale preparation of the oligomeric peptides. To date, research on *Nostoc* symbioses has focused predominantly on diffusible chemical signals that regulate motility, chemoattraction, and cellular differentiation. ^2,3,14^ Our findings suggest that contact-dependent interactions represent an additional and previously underappreciated layer of symbiotic communication. Such interactions may, for example, influence whether *Nostoc* establishes predominantly epiphytic associations, as observed in feather mosses, or endophytic associations, as in *Blasia*.^15,16^ Collectively, our results uncover a previously unrecognized mode of lanthipeptide maturation through covalent oligomerization and identify nostolanthin as a host-responsive RiPP that promotes the establishment of physical contact during the early transition from a free-living to a symbiotic lifestyle.

## Methods

### Cultivation of cyanobacteria

Wild-type *N. punctiforme* cultures were maintained in liquid BG11 ^39^ medium supplemented with 5 mM NaHCO_3_, wild-type *Nostoc* sp. KVJ2 cultures were maintained in liquid BG11_0_ medium ^39^ under continuous white light with 20 µmol photons m^-2^s^-1^ at 22 °C. Mutant strains were cultivated in BG11 medium supplemented with 5 mM NaHCO_3_ and 2 µg mL^-1^ streptomycin under continuous white light with 18 µmol photons m^-2^s^-1^ at 25 °C.

### Cultivation of *Blasia pusilla*

*B. pusilla* is maintained in Erlenmeyer flasks containing regeneration medium (1/5 BG11 supplemented with 6.25 mM NaNO₃) under continuous white light illumination with 30 µmol photons m^-2^s^-1^ at 22 °C. To generate nitrogen-starved liverwort for symbiosis assays, *B. pusilla* is cultivated for 5 weeks in nitrogen-starvation medium (1/5 BG11_0_).

### N. punctiforme - B. pusilla interaction studies

Conditioned *B. pusilla* medium was prepared by filtering the culture supernatant, transferring it into 50 mL Falcon tubes, and concentrating it 100-fold to a final volume of 500 µL using a vacuum concentrator (RVC 2-25 CDplus, Christ, Germany; 1 mbar, 30 °C, 1400 rpm, 15 h). For chemical interaction assays, concentrated *B. pusilla* exudate was added to *N. punctiforme* wild-type or reporter strain cultures to achieve the equivalent of 50% (v/v) unconcentrated exudate (e.g., 7.5 µL of 100× concentrate added to 15 mL culture). For physical interaction studies, nitrogen-deprived moss was cut into ∼1 cm² pieces (approximately 1 g wet weight) and incubated overnight in fresh 1/5 BG11_0_ medium. For co-cultivation, 15 mL of *N. punctiforme* culture (wild-type or reporter strain) was combined with one *B. pusilla* segment in a 25 mL Erlenmeyer flask. All conditions were prepared in triplicate, including no-moss controls as references. Co-cultures were sampled at defined time points and processed either for RNA extraction or confocal fluorescence microscopy.

### RNA isolation and RT-PCR

Cells with a wet weight of approximately 0.5 g were pelleted; RNA was isolated by using the hot-TRIzol method (Life Technologies GmbH, Darmstadt, Germany), and supernatants were purified with an RNeasy kit, including on-column DNase digestion (Qiagen, Hilden, Germany). First-strand reverse transcription was carried out using Maxima reverse transcriptase with random oligonucleotides (Thermo Fisher Scientific). For RT-qPCR, a LightCycler 480 (Roche Applied Science, Mannheim, Germany) in combination with a Sybr green-based detection system (SensiFAST SYBR Lo-ROX kit; Bioline, Luckenwalde, Germany) was used. Specific primer pairs for individual BGCs were designed previously. The RNase P encoding gene *rnpB* was used as a housekeeping gene for standardization, as described previously (11). All primers were tested in PCRs prior to RT-qPCR. Each reaction was carried out in four technical replicates. Raw data were converted using the software LC480Converter; subsequent processing and calculation according to the Pfaffl method were carried using LinRegPCR software. ^40^

### Fluorescence microscopy

Fluorescence microscopy was performed with a confocal laser scanning microscope (LSM 710; Carl Zeiss, Jena, Germany). Images were recorded with filter presets for chlorophyll a and enhanced CFP (eCFP). For device control, image acquisition and processing the ZEN software were used. Care was taken to keep all image recording parameters constant for all samples.

### Construction and site-directed mutagenesis of *E. coli* expression vectors

Genes encoding the precursor peptide 1 (RS25510) and the lanthipeptide synthetase (RS25520) were amplified by PCR with Q5 polymerase from genomic DNA of *N. punctiforme* using primers NB46/NB47 and NB16/NB17, respectively. Amplified fragments were gel-purified (GeneJET Gel Extraction Kit, Thermo Fisher Scientific). The precursor peptide gene was inserted into pET28a (linearized with NdeI) to generate an N-terminally His-tagged construct, and the synthetase gene was inserted into pCDFDuet-1 (linearized with NcoI) to generate an untagged construct for co-expression. Linearized vectors were gel-purified (GeneJET Gel Extraction Kit, Thermo Fisher Scientific), and inserts were incorporated using HiFi DNA Assembly (NEB). Four site-directed variants of the precursor peptide gene were generated by Q5 site-directed mutagenesis (NEB) using the primers listed in Table S3B. All plasmids were transformed into chemically competent *E. coli* XL1-Blue cells (Agilent Technologies) and verified by Sanger sequencing (LGC, Biosearch Technologies).

### Peptide expression

For heterologous expression, constructs carrying the genes encoding the N-terminally His-tagged precursor peptide and the lanthionine synthetase were co-transformed into *E*. *coli* LOBSTR. ^41^ A single transformed colony was selected from a lysogeny broth (LB) agar plate and used to inoculate into LB broth supplemented with kanamycin (40 μg/mL) and streptomycin (20 μg/mL). The culture was incubated overnight at 37 °C and 180 rpm. The overnight culture was then used to inoculate baffled Erlenmeyer flasks containing terrific broth supplemented with the same antibiotic. Expression cultures were incubated at 37 °C and 200 rpm until reaching an optical density measured at 600 nm (OD600) of 1.0. Peptide expression was induced by adding 0.5 mM (final concentration) isopropyl β-D-thiogalactopyranoside (IPTG), followed by incubation at 18 °C and 200 rpm for 18 h. Cells were harvested by centrifugation (10 min, 4°C, 6000 × g), and the resulting pellets were stored at-20 °C until further purification. The N-terminally His-tagged peptidase LahT150 was expressed under identical conditions, except that LB broth supplemented with carbenicillin (100 μg/mL) was used.

### Purification of precursor peptides under native conditions

All steps for peptide purification were performed at 4 °C or on ice. Cells containing the expressed His_6_-precursor peptide were thawed, resuspended in a peptide lysis buffer (50 mM NaH_2_PO_4_, 300 mM NaCl, pH 8.0) and lysed by sonication (Sonopuls HD ultrasonic homogenizer, 10 min, 70 % amplitude, pulse 3 s on/off). To the lysed cells, imidazole was added for a final concentration of 10 mM and the cells were centrifuged (20 min, 4 °C, 10000 × g). The supernatant was incubated with 1 mL PureCube Ni-NTA agarose at 4 °C for 1 h. Subsequently, the resin was washed twice with 5 mL peptide lysis buffer containing 20 mM imidazole. The peptide was eluted twice with 1 mL peptide lysis buffer containing 500 mM imidazole. The peptide was concentrated using Amicon Ultra-4 centrifugal filters (3 kDa MWCO, Merck Millipore).

### Purification of precursor peptide under denaturing conditions

Cell pellets containing the overexpressed precursor His_6_-NltA1 from *N. punctiforme* were thawed on ice and resuspended in denaturing lysis buffer (100 mM NaH_2_PO_4_, 10 mM Tris, 8 M urea, pH 8.0) at 5-10 mL g^-^¹ wet weight. Cells were lysed by end-over-end shaking (30-60 min, room temperature) combined with gentle vortexing until the lysate became translucent, and cellular debris was removed by centrifugation (10,000 × g, 20 min, 22 °C). The soluble fraction was incubated with Ni-NTA resin (pre-equilibrated in denaturing lysis buffer) for 1 h at room temperature with end-over-end mixing. The resin was pelleted (4,700 × g, 5 min, 22 °C) and washed 3-4 times with denaturing wash buffer (100 mM NaH_2_PO_4_, 10 mM Tris, 8 M urea, pH 6.3). The His-tagged protein was eluted five times with denaturing elution buffer (100 mM NaH_2_PO_4_, 10 mM Tris, 8 M urea, pH 4.5), with each eluate collected separately. Owing to urea dissociation, all buffers were pH-adjusted immediately before use.

### Purification of leader peptidase LahT150

The peptidase LahT150 was purified following the same protocol as described for the precursor peptides, with the exception that the LahT150 lysis buffer contained 20 mM Tris and 1 M NaCl (pH 7.8). The concentrated protein was subsequently subjected to size-exclusion chromatography on a HiLoad 16/600 Superdex 200 column (GE Healthcare), eluting with 1.5 column volumes (CV) of LahT150 storage buffer (20 mM Tris, 1 M NaCl, 10% glycerol, pH 7.8), as previously described by Bobeica et al. ^42^

### Proteolytic digests of precursor peptides

Leader peptide cleavage was carried out by incubating 500 µM peptide with 100 µM of the peptidase LahT150 in 100 mM Tris (pH 8.0) supplemented with 1 mM TCEP at 37 °C for 3 h.

### Iodoacetamide (IAA) assay

For the IAA assay, 2 µL IAA (500 mM), 1 µL TCEP (50 mM), and 27 µL Tris buffer (pH 8.6) were added to 20 µL of the LahT150 digestion reaction. The mixture was incubated for 4 h at room temperature in the dark. The 50 µL assay was then diluted with 60 µL H₂O (LC-MS grade) and analyzed by LC-MS.

### SDS-PAGE and western blot preparation

Prior to SDS-PAGE and western blot analysis, samples were mixed with 5× SDS loading buffer (250 mM Tris, pH 6.8, 10% (w/v) SDS, 50% (v/v) glycerol, 0.1% (w/v) bromophenol blue, 500 mM 2-mercaptoethanol) and boiled at 95 °C for 10 min.

### HPLC purification

After IMAC purification, the modified and LahT150-cleaved peptides were further purified by RP-HPLC on a Thermo Fisher Vanquish HPLC system equipped with a Hypersil GOLD™ C18 column (5 µm, 250 mm × 4.6 mm) maintained at 40 °C, using a linear gradient from 5% to 95% solvent B (100% ACN, 0.1% TFA, LC-MS grade) in solvent A (water, 0.1% TFA, LC-MS grade) at a flow rate of 1 mL/min over 20 min. For purification of nostolanthin-T17A for NMR analysis, the gradient was extended to 95% solvent B over 30 min. Absorbance was monitored at 218 nm. Fractions containing the cleaved, modified core peptide were dried using a SpeedVac and stored at-20 °C until further characterization by LC-MS or NMR.

### Mass spectrometric analysis of peptides

Analytical HPLC-HRMS was performed on a Thermo Scientific Vanquish Flex LC system coupled to an Orbitrap Exploris 240 mass spectrometer. The samples (5 µL injections) were separated on a Phenomenex Kinetex® C18 (50 × 2.1 mm, 1.7 µm particle size, 100 Å pore size) heated to 40 °C. HPLC separation was performed with the following standard methods (solvent A: H2O +0.1% FA; solvent B: acetonitrile (ACN) +0.1% FA, flow rate: 0.5 mL/min): 1 min at 5% B; 1–11 min from 5% to 100% B; 11–13 min at 100% B; 13–14 min from 100% to 5%. The mass spectrometer was run in positive ion mode with heated electrospray ionization (spray voltage: 3500 V, ion transfer tube temperature: 300 °C, vaporizer temperature: 350 °C, S-lens RF level 70%). Full MS scans were acquired over an m/z range of 250-2000 at a resolution of 30,000 (AGC target: 2e5, maximum injection time: 54 ms). MS2 fragmentation was performed at a resolution of 15,000 (AGC target: 2e4, maximum injection time: 54 ms, isolation window: 1.0 m/z) with a normalized collision energy (NCE) of 30.

### NMR spectroscopy

Lyophilized nostolanthin-T17A (1.3 mg) purified from *E. coli* was dissolved in 600 µL of an H_2_O:D_2_O mixture (9:1) (Deutero GmbH, Kastellaun, Germany), and filled into a 5-mm NMR sample tube. NMR experiments were performed on a Bruker Avance III 700 MHz spectrometer equipped with a room temperature TXI probe (Karlsruhe, Germany). ^1^H chemical shifts were referenced internally with trimethylsilylpropanoic acid (TMSP-*d*4, Deutero GmbH, Kastellaun, Germany). ^13^C chemical shifts were then referenced with respect to ^1^H using a correction factor of *f*_13C/1H_ = 0.251449530. ^43,44^

After initial tests at different temperatures, a full data set was recorded at 308 K including 1D ^1^H, 2D ^1^H-^1^H TOCSY, ^1^H-^1^H NOESY and ^1^H-^13^C HSQC spectra. A second data set was recorded for nostolanthin-T17A in 100% D_2_O after evaporation of the protonated solvent.

Homonuclear 2D spectra were recorded with acquisition times of 160 ms and 30 ms in the direct and indirect ^1^H dimension, respectively. For ^1^H-^1^H TOCSY experiments, an MLEV17 sequence with a radio-frequency field strength of 10 kHz and a duration of 60 ms was used for the homonuclear Hartmann-Hahn transfer. ^1^H-^1^H NOESY spectra were recorded with a mixing time of 300 ms. ^1^H-^13^C HSQC spectra were recorded with acquisition times of 160 ms and 9 ms in the direct ^1^H and indirect ^13^C dimension, respectively. Non-uniform sampling (50% sparse sampling) was employed for TOCSY and HSQC spectra.

2D NMR data were processed by applying linear forward prediction and zero filling prior to Fourier transformation. Apodization of time domain data was performed using a squared sine bell function shifted by 90-180°. Data acquisition and processing were performed via TopSpin 3.1 (Bruker, Germany), whereas POKY was used for resonance assignment and data visualization. ^45^

### Generation of the anti-NltA1 antibody

To generate a nostolanthin-specific antibody, the precursor peptide co-expressed with NltM in *E. coli* was purified under native conditions as described above. Following concentration, the sample was incubated in carbonate-bicarbonate buffer (12.5 mM NaHCO_3_, 87.5 mM Na_2_CO_3_, pH 10.6) for 48 h at room temperature to promote oligomerization prior to immunization. The sample was clarified by filtration through a 0.45 µm cellulose acetate filter and applied to a Superdex 200 Increase 10/300 GL column (bed volume: 24 mL) pre-equilibrated with carbonate-bicarbonate buffer, using an ÄKTA pure system (Cytiva) at a flow rate of 0.75 mL/min under isocratic conditions at room temperature. Elution was monitored by UV absorbance at 280 nm, and fractions were collected accordingly. The fraction eluting at 8.86 mL (Supplementary Fig. 16) was concentrated using Amicon Ultra-4 centrifugal filters (30 kDa MWCO, Merck Millipore) and submitted to BioGenes (Berlin, Germany) as an antigen (3 mg) for generation of a nostolanthin-specific antibody. Immunization was performed in a rabbit (animal RB02263; order no. 59647, lot no. 99181) according to a standard immunization protocol comprising an initial immunization followed by five boosts over a 13-week period, with intermediate and final bleedings collected throughout (Table S6). Antiserum titer, determined after the third boost, was reported as 1:200,000. The final antiserum was stored at 4 °C and used crude for subsequent applications. The antibody was validated by testing against the nostolanthin precursor monomer and oligomer, both purified from *E. coli* after co-expression with NltM and separated by SEC (Supplementary Fig. 16 and 18). The polyclonal anti-NltA1 antibody recognized both the precursor monomer and oligomer. For Dot-blots, the anti-NltA1 antibody was diluted 1:20,000 in TBS-T.

### Generation of *Nostoc* overexpression mutants

For the generation of the pRL1049-nltA1vector, the previously constructed plasmid pJK008 was used which contains a streptomycin resistance cassette and a strong constitutive promoter (5’ UTR of *RS16340*) from *N. punctiforme*, followed by a downstream *EcoRI* restriction site. ^29^ Primers NB37/NB38 were used to amplify the modified, His-tagged precursor peptide gene *RS25510* from a previously synthesized DNA template. ^46^ The precursor peptide gene was introduced into pJK008 using HiFi-Builder (NEB) mediated homologous recombination and the construct was transformed into chemically competent *E. coli* XL-1 blue cells (Agilent Technologies, Waldbronn, Germany) by transformation. Prior to transformation into *N. punctiforme* the construct was verified by sequencing (LGC, Biosearch Technologies).

To construct the vector pRL1049-nlt_BGC, the first half of the *nlt* BGC (*allorf_6250196_6250426* - *RS25520*) was amplified using primers NB85/NB86 and genomic DNA of *N. punctiforme* as a template. It was integrated into the *EcoRI* restriction site of pJK008 using HiFi-Builder, incorporating a *XhoI* restriction site downstream of *RS25520*. The resulting construct was transformed into chemically competent *E. coli* XL-1 blue cells by transformation and verified by sequencing. Then the second half of the BGC (upstream intergenic region of *RS25525* - *RS25545*), amplified using primers NB88/NB89 and genomic DNA of *N. punctiforme* as a template was then introduced into the *XhoI* site of the previously constructed vector using HiFi-Builder. The final construct was transformed into chemically competent *E. coli* XL-1 blue cells by transformation and verified by sequencing.

For transformation of *Nostoc* cultures, a dense culture was harvested, washed four times with sterile water, and concentrated to a final volume of 400 µL sterile water. The overexpression plasmid (10 µg) was added to the cell suspension and introduced by electroporation (Micropulser™, Bio-Rad, Munich, Germany; 4 ms, 1.5 kV, 1 pulse). Cells were plated on BG11 medium supplemented with 5 mM NaHCO_3_, overlaid with GSTF mixed cellulose ester filters, and incubated under low light (10 µmol photons m^-2^s^-1^) at 22 °C for 2 days. After a further 5 days under standard light conditions (30 µmol photons m^-2^s^-1^), filters were transferred onto selective BG11 plates containing 5 mM NaHCO_3_ and 2 µg mL^-1^ streptomycin. Non-resistant background cells died within 2-3 weeks. Resistant colonies were subsequently transferred into liquid medium. Successful transformation was confirmed by colony PCR.

### Purification of KVJ2-*nlt* supernatant

The culture supernatant of *Nostoc* sp. KVJ2-*nlt* was freeze-dried and reconstituted in protein buffer (100 mM Tris, 150 mM NaCl, 10% (w/w) glycerol, pH 7.5). The sample was clarified by filtration through a 0.45 µm cellulose acetate filter and applied to a Superdex 200 Increase 10/300 GL column (bed volume: 24 mL) pre-equilibrated with protein buffer, using an ÄKTA pure system (Cytiva) at a flow rate of 0.75 mL/min under isocratic conditions at room temperature. Elution was monitored by UV absorbance at 280 nm, and fractions were collected accordingly. Fractions were concentrated using Amicon Ultra-4 centrifugal filters (Merck Millipore) and stored at 4 °C until further use.

### Nostoc - B. pusilla symbiosis assay

Nitrogen-starved *B. pusilla* was cut into approx. 2 × 1 cm sections and transferred into 50 mL flasks containing fresh BG11_0_ medium. A dense culture of *Nostoc* sp. KVJ2 was washed with BG11_0_, and equal wet biomass was added to three flasks. Nostolanthin monomer (0.1 mg) was added to one flask, nostolanthin oligomer (20 µL of the SEC-fraction) to a second flask, and the third flask served as an untreated control. Flasks were incubated at 55 rpm and 26 °C under standard low-light conditions (23 µmol photons m^-2^s^-1^)

## Supporting information

complete SI section

## Acknowledgements

We are grateful to Anja Schüler for technical assistance. This study was supported by the Deutsche Forschungsgemeinschaft (DFG, German Research Foundation) as part of the DFG-funded graduate school RTG 2473 (Bioactive Peptides, project number 392923329) to N.B., E.D. and R.D.S. The mass spectrometer used in this study was partially funded by the DFG (project number 467315902) to E.D.

