## Supplementary material for "An Oligomeric Lanthipeptide from *Nostoc punctiforme* Promotes Host Association During Early Symbiosis with *Blasia pusilla*": complete SI section

#### Content

#### 1. Supplementary tables

**Table S1:** Vectors used in this study.
**Table S2:** Strains used in this study.
**Table S3:** Primers used in this study. A) Primers used for Gibson Assembly. B) Primers used for site-directed mutagenesis. C) Primers used for RT-qPCR.
**Table S4:** Functional annotation of genes within the nostolanthin BGC.
**Table S5:** $^{1}H$ and $^{13}C$ resonance assignments of nostolanthin-T17A recorded in $H_2O$ at 308 K.
**Table S6:** Immunization schedule for generation of the anti-NltA1 polyclonal antibody.

#### 2. Supplementary figures

**Figure S1.** HPLC profiles of cell extracts of *B. pusilla* and *N. punctiforme* either grown in mono-culture of co-culture for 6h.
**Figure S2.** Representative BGCs closely related to the *nlt* BGC based on a BLASTP search of the precursor peptides, the lanthipeptide synthetase and the response regulators.
**Figure S3.** LC-MS analysis of LahT150-cleaved $His_6$-NltA1 co-expressed with NltM in *E. coli* identifies Nostolanthin A and a glutathionylated adduct.
**Figure S4.** LC-MS and LC-MS/MS analysis of LahT150-cleaved $His_6$-NltA1-S5A co-expressed with NltM in *E. coli*.
**Figure S5.** Iodoacetamide (IAA) assay of LahT150-cleaved $His_6$-NltA1-S5A co-expressed with NltM in *E. coli*.
**Figure S6.** LC-MS and LC-MS/MS analysis of LahT150-cleaved $His_6$-NltA1-C11A co-expressed with NltM in *E. coli*.
**Figure S7.** Iodoacetamide (IAA) assay of LahT150-cleaved $His_6$-NltA1-C11A co-expressed with NltM in *E. coli*.
**Figure S8.** LC-MS and LC-MS/MS analysis of LahT150-cleaved $His_6$-NltA1-C12A co-expressed with NltM in *E. coli*.
**Figure S9.** Iodoacetamide (IAA) assay of LahT150-cleaved $His_6$-NltA1-C12A co-expressed with NltM in *E. coli*.
**Figure S10.** LC-MS and LC-MS/MS analysis of LahT150-cleaved $His_6$-NltA1-T17A co-expressed with NltM in *E. coli*.
**Figure S11.** Iodoacetamide (IAA) assay of LahT150-cleaved $His_6$-NltA1-T17A co-expressed with NltM in *E. coli*.
**Figure S12.** Amide region of the 1H-1H TOCSY spectrum of nostolanthin-T17A recorded in H2O at 308 K.
**Figure S13.** Overlay of the amide regions of $^{1}H$-$^{1}H$ TOCSY (magenta) and $^{1}H$-$^{1}H$ NOESY (cyan) spectra of nostolanthin-T17A recorded in $H_2O$ at 308 K.
**Figure S14.** $^{1}H$-$^{13}C$ HSQC spectrum of nostolanthin-T17A recorded in $H_2O$ at 308 K.
**Figure S15.** Bioactivity assay with nostolanthin A.
**Figure S16.** Analytical size-exclusion chromatography (SEC) analysis of NltA1.
**Figure S17.** RT-qPCR of the *nlt* BGC in *Nostoc* sp. KVJ2-*nlt*.
**Figure S18.** Western blot to test the polyclonal antibody anti-NltA1.

#### 3. Supplementary references

### 1. Supplementary tables

**Table S1**. Vectors used in this study.

| Name | Source | Resistance marker |
|---|---|---|
| pET28a | Sigma-Aldrich, St. Louis, MO, USA | Kanamycin |
| pET28a-nltA1 | This work | Kanamycin |
| pET28a-nltA1_S5A | This work | Kanamycin |
| pET28a-nltA1_C11A | This work | Kanamycin |
| pET28a-nltA1_C12A | This work | Kanamycin |
| pET28a-nltA1_T17A | This work | Kanamycin |
| pCDFDuet-1 | Novagene | Streptomycin |
| pCDF-nltM | This work | Streptomycin |
| pETDuet-LahT150 | (1) | Ampicillin |
| pRL1049 | (2) | Streptomycin |
| pRL1049-nltA1 | This work | Streptomycin |
| pRL1049-nlt BGC | This work | Streptomycin |

**Table S2**. Strains used in this study.

| Organism | Phylum | Source |
|---|---|---|
| *E. coli* XL1-blue | proteobacteria | Agilent Technologies, Waldbronn, Germany |
| *E. coli* LOBSTR | proteobacteria | Kerafast, Inc., Boston, MA, USA |
| *Nostoc punctiforme* PCC 73102 | cyanobacteria | Pasteur Culture Collection |
| *Nostoc* sp. KVJ2 | cyanobacteria | (3) |

**Table S3**. Primers used in this study. A) Primers used for Gibson Assembly. B) Primers used for site-directed mutagenesis. Lowercase letters indicate the mutated nucleotides. C) Primers used for RT-qPCR.

A)

| Construct | Name | Sequence (5' to 3') | Target |
|---|---|---|---|
| pET28a-nltA1 | NB46 _fw | CTGGTGCCGCGCGGCAGCCATATGTTACACCAAAT CAAAG | nltA1 (NPUN_RS25510) |
| pET28a-nltA1 | NB47 _rv | TCCACCAGTCATGCTAGCCATTATCTCGTTGGACAG TC | nltA1 (NPUN_RS25510) |
| pCDF-nltM | NB16_fw | ACTTTAATAAGGAGATATACATGTCTCAACTATTTGT TCAATC | nltM (NPUN_RS25520) |
| pCDF-nltM | NB17_rv | TGATGGTGATGGCTGCTGCCTCATTCCCATAACAGC AC | nltM (NPUN_RS25520) |
| pRL1049-nltA1 | NB37_fw | AATCCAATAGGAGAAAAAGGATGAGAGGATCGCATC | nltA1 (NPUN_RS25510) |
| pRL1049-nltA1 | NB38_rv | GAGGCCCTTTCGTCTTCAAGTTATCTCGTTGGACAG TC | nltA1 (NPUN_RS25510) |
| pRL1049-nlt_BGC | NB85_fw | AATCCAATAGGAGAAAAAGGTTGGCGTTGCTGAATT C | nlt BGC_1 (allorf_6250196_6250426) |
| pRL1049-nlt_BGC | NB86_rv | GAGGCCCTTTCGTCTTCAAGCTCGAGTCATTCCCAT AACAGCAC | nlt BGC_2 (NPUN_RS25520) |
| pRL1049-nlt_BGC | NB88_fw | CGGTGCTGTTATGGGAATGACCTAGAAGTCATCAAA ATATTCTC | nlt BGC_3 (NPUN_RS25520-NPUN_RS25525) |
| pRL1049-nlt_BGC | NB89_rv | GAGGCCCTTTCGTCTTCAAGCTTATGACTGCCGAAC TAC | nlt BGC_4 (NPUN_RS25545) |

B)

| Name | Sequence (5' to 3') |
|---|---|
| NB190_RiPP4_S5A_fw_NEB | CGTCTCTGAAgcaCACGTACACA |
| NB191_RiPP4_S5A_rv_NEB | GCTCCTCCACTAACGC |
| NB194_RiPP4_C11A_fw_NEB | ACACATGTCAgctTGTAACGACTGTCC |
| NB195_RiPP4_C11A_rv_NEB | ACGTGTGATTCAGAGAC |
| NB196_RiPP4_C12A_fw_NEB | CATGTCATGTgctAACGACTGTCCAAC |
| NB197_RiPP4_C12A_rv_NEB | TGTACGTGTGATTCAG |
| NB200_RiPP4_T17A_fw_NEB | CGACTGTCCAgcgAGATAATGGC |
| NB201_RiPP4_T17A_rv_NEB | TTACAACATGACATGTGTACGTG |

C)

| Name | Sequence (5' to 3') | Target | Amplicon size [bp] |
|---|---|---|---|
| C3_FW | CGTGGTTGACTGGAGATGCT | Npun_RS10475, PKS1 (nvl/ncl) | 108 |
| C3_RV | AAGCCTCTCGTGCCGTTTTA | Npun_RS10475, PKS1 (nvl/ncl) | 108 |
| D132_FW | GCGGACTAGCTCATCAGACC | Npun_RS15970, PKS2 | 118 |
| D133_RV | GTACCATCCCCACAACCCAT | Npun_RS15970, PKS2 | 118 |
| D136_FW | AGAACGGGCGCTACTCTTTT | Npun_RS17005, PKS3 (ngl) | 105 |
| D137_RV | TTTCTTGGTGGATAGGCGGG | Npun_RS17005, PKS3 (ngl) | 105 |
| D138_FW | ACTCTCAGGCGAATGTTCCA | Npun_RS17455, PKS4 | 144 |
| D139_RV | CCTGAAATTGACGCGCAGAT | Npun_RS17455, PKS4 | 144 |
| C21_FW | GGGGAATGGAAAGCATGGGA | Npun_RS33540, PKS5 | 125 |
| C21_RV | ATTAACGCCCCTTCCCTGTG | Npun_RS33540, PKS5 | 125 |
| C8_FW | AGCGGCAACATATTCCCCAA | Npun_RS42520, NRPS1 | 113 |
| C8_RV | ACGCCCAACCTGCTCTATTC | Npun_RS42520, NRPS1 | 113 |
| Cp_FW | CCTCAATCCAAGTCAGGCGT | Npun_RS38145, NRPS2 | 136 |
| Cp_RV | CTCATGTCGGGTGCAGCTTA | Npun_RS38145, NRPS2 | 136 |
| apt_RT_FW | GAAATTGAGGCGCTTTTGAG | Npun_RS12350, Apt | 136 |
| apt_RT_RV | GGCTAGTGACGCTCACATCA | Npun_RS12350, Apt | 136 |
| NosA_RT_FW | GTTTGCCCTCTCTGCTGAAC | Npun_RS10970, NosA | 109 |
| NosA_RT_RV | GCGGTAAAGCAGGGTATCAA | Npun_RS10970, NosA | 109 |
| C10L_FW | GGCAGAATTGGGAGGACGAA | Npun_RS16255, RiPP1a | 111 |
| C10L_RV | TCCCAAACCCATCATTGAGCA | Npun_RS16255, RiPP1a | 111 |
| D134_FW | CGAAAGAAGCAGTCATCAAA | Npun_RS16310, RiPP1b | 164 |
| D135_RV | TTGCTTTTCTTGCATCACTG | Npun_RS16310, RiPP1b | 164 |
| C11a_FW | AGCAGACATCATAGCTCCACT | Npun_RS16795, RiPP3 | 130 |
| C11a_RV | GGGTGCAGAAAAGGGCTACA | Npun_RS16795, RiPP3 | 130 |
| D140_FW | GCGATCTCTCAAATGCTGGC | Npun_RS25510, RiPP4 (nlt) | 103 |
| D141_RV | CGTGTGATTCAGAGACGGCT | Npun_RS25510, RiPP4 (nlt) | 103 |
| D150_FW | GCTAGGAGCTACTGAGAACA | Npun_AF077, RiPP5 (npu) | 97 |
| D151_RV | AGCATCTACCTCCTGAATCG | Npun_AF077, RiPP5 (npu) | 97 |
| rnpB_RT3_FW | GCGGTTGCAGATCAGTCATA | Npun_R018, RnpB | 110 |
| rnpB_RT3_RV | TCTGTGGCACTATCCTCACG | Npun_R018, RnpB | 110 |

**Table S4** Functional annotation of genes within the nostolanthin BGC.

| Gene | Amino acids | Proposed function | Accession number |
|---|---|---|---|
| nltR1 | 511 | Hybrid sensor histidine kinase/response regulator, PAS domain | WP_012411336.1 |
| nltR2 | 124 | response regulator, DNA-binding receiver domain | WP_012411337.1 |
| nltA1 | 91 | Nif11-like leader peptide family RiPP precursor | WP_012411338.1 |
| nltA2 | 91 | Nif11-like leader peptide family RiPP precursor | WP_012411339.1 |
| nltM | 1101 | type 2 lanthipeptide synthetase LanM family protein | WP_012411340.1 |
| nltT1 | 479 | ABC transporter, NHLP family, membrane fusion subunit | WP_012411341.1 |
| nltT2 | 723 | ABC transporter, NHLP family, permease subunit, C39 peptidase domain | WP_234710978.1 |
| nltT3 | 961 | ABC transporter, NHLP family, ATP-binding subunit | WP_012411344.1 |

**Table S5** $^{1}H$ and $^{13}C$ resonance assignments of nostolanthin-T17A recorded in $H_2O$ at 308 K.

| Residue | Atom | δ (ppm) | sd (ppm) | Number of assignments |
|---|---|---|---|---|
| **Ala1** | CB | 19.557 | 0.000 | 1 |
| | HA | 4.189 | 0.006 | 3 |
| | HB | 1.552 | 0.002 | 3 |
| **Val2** | CB | 32.971 | 0.000 | 1 |
| | CG1 | 21.164 | 0.000 | 1 |
| | CG2 | 20.562 | 0.000 | 1 |
| | HA | 4.223 | 0.007 | 4 |
| | HB | 2.097 | 0.008 | 5 |
| | HG1 | 0.970 | 0.005 | 3 |
| | HG2 | 0.972 | 0.005 | 5 |
| | HN | 8.490 | 0.002 | 6 |
| **Ser3** | CB | 63.971 | 0.001 | 2 |
| | HA | 4.521 | 0.006 | 4 |
| | HB2 | 3.923 | 0.007 | 4 |
| | HB3 | 3.884 | 0.005 | 4 |
| | HN | 8.460 | 0.003 | 6 |
| **Glu4** | CB | 32.785 | 0.000 | 1 |
| | CG | 28.819 | 0.024 | 2 |
| | HA | 4.379 | 0.004 | 6 |
| | HB23 | 2.538 | 0.004 | 5 |
| | HG2 | 2.169 | 0.004 | 5 |
| | HG3 | 2.056 | 0.007 | 5 |
| | HN | 8.524 | 0.001 | 5 |
| **Lan5** | CB | 36.410 | 0.009 | 2 |
| | HA | 4.319 | 0.005 | 6 |
| | HB2 | 3.204 | 0.009 | 7 |
| | HB3 | 2.958 | 0.011 | 6 |
| | HN | 8.693 | 0.000 | 6 |
| **His6** | CB | 28.447 | 0.008 | 2 |
| | CD2 | 120.026 | 0.000 | 1 |
| | CE1 | 136.632 | 0.000 | 1 |
| | HA | 4.599 | 0.007 | 4 |
| | HB2 | 3.383 | 0.007 | 5 |
| | HB3 | 3.279 | 0.004 | 5 |
| | HD2 | 7.300 | 0.006 | 4 |
| | HE1 | 8.675 | 0.008 | 2 |
| | HN | 8.474 | 0.003 | 7 |
| **Val7** | CB | 32.847 | 0.000 | 1 |
| | CG1 | 20.849 | 0.000 | 1 |
| | CG2 | 21.266 | 0.000 | 1 |

| | | | | |
|---|---|---|---|---|
| | HA | 4.065 | 0.005 | 5 |
| | HB | 2.060 | 0.006 | 7 |
| | HG1 | 0.858 | 0.003 | 10 |
| | HG2 | 0.807 | 0.007 | 6 |
| | HN | 7.795 | 0.004 | 8 |
| **His8** | CB | 28.313 | 0.010 | 2 |
| | CD2 | 120.158 | 0.000 | 1 |
| | CE1 | 136.627 | 0.000 | 1 |
| | HA | 4.780 | 0.006 | 4 |
| | HB2 | 3.340 | 0.010 | 7 |
| | HB3 | 3.219 | 0.004 | 6 |
| | HD2 | 7.330 | 0.003 | 6 |
| | HE1 | 8.665 | 0.000 | 1 |
| | HN | 8.206 | 0.003 | 7 |
| **Met9** | CB | 33.569 | 0.014 | 2 |
| | CG | 32.197 | 0.000 | 1 |
| | CE | 17.040 | 0.000 | 1 |
| | HA | 4.555 | 0.008 | 5 |
| | HB2 | 2.154 | 0.004 | 4 |
| | HB3 | 2.053 | 0.004 | 4 |
| | HG23 | 2.626 | 0.005 | 5 |
| | HE | 2.135 | 0.000 | 1 |
| | HN | 8.203 | 0.001 | 5 |
| **Lan10** | CB | 36.886 | 0.014 | 2 |
| | HA | 4.717 | 0.007 | 3 |
| | HB2 | 3.420 | 0.005 | 6 |
| | HB3 | 3.189 | 0.006 | 4 |
| | HN | 8.853 | 0.002 | 4 |
| **Cys11** | CB | 27.599 | 0.000 | 1 |
| | HA | 4.569 | 0.009 | 3 |
| | HB23 | 3.103 | 0.006 | 6 |
| | HN | 8.206 | 0.001 | 2 |
| **Lan12** | CB | 37.061 | 0.002 | 2 |
| | HA | 4.743 | 0.009 | 4 |
| | HB2 | 3.255 | 0.006 | 6 |
| | HB3 | 3.005 | 0.007 | 6 |
| | HN | 8.056 | 0.002 | 7 |
| **Asn13** | CB | 37.779 | 0.007 | 2 |
| | HA | 4.500 | 0.006 | 5 |
| | HB2 | 2.975 | 0.009 | 4 |
| | HB3 | 2.858 | 0.006 | 7 |
| | HD1 | 7.568 | 0.002 | 2 |
| | HD2 | 6.882 | 0.011 | 2 |

| | | | | |
|---|---|---|---|---|
| | HN | 8.613 | 0.002 | 8 |
| **Asp14** | CB | 37.717 | 0.000 | 1 |
| | HA | 4.795 | 0.002 | 4 |
| | HB23 | 2.893 | 0.006 | 5 |
| | HN | 7.948 | 0.003 | 7 |
| **Lan15** | CB | 36.358 | 0.017 | 2 |
| | HA | 4.855 | 0.012 | 6 |
| | HB2 | 3.181 | 0.006 | 5 |
| | HB3 | 3.043 | 0.008 | 5 |
| | HN | 7.984 | 0.002 | 6 |
| **Pro16** | CB | 32.104 | 0.031 | 2 |
| | CG | 27.402 | 0.001 | 1 |
| | CD | 50.832 | 0.006 | 2 |
| | HA | 4.454 | 0.004 | 7 |
| | HB2 | 2.323 | 0.005 | 7 |
| | HB3 | 1.974 | 0.004 | 6 |
| | HG23 | 2.044 | 0.003 | 6 |
| | HD2 | 3.810 | 0.005 | 7 |
| | HD3 | 3.691 | 0.004 | 9 |
| **Ala17** | CB | 19.299 | 0.000 | 1 |
| | HA | 4.338 | 0.002 | 3 |
| | HB | 1.427 | 0.004 | 4 |
| | HN | 8.311 | 0.001 | 5 |
| **Arg18** | CB | 30.832 | 0.019 | 2 |
| | CG | 27.146 | 0.000 | 1 |
| | CD | 43.459 | 0.000 | 1 |
| | HA | 4.379 | 0.002 | 5 |
| | HB2 | 1.957 | 0.005 | 5 |
| | HB3 | 1.811 | 0.006 | 5 |
| | HG23 | 1.674 | 0.006 | 5 |
| | HD23 | 3.250 | 0.004 | 7 |
| | HE | 7.193 | 0.004 | 4 |
| | HN | 8.271 | 0.001 | 6 |

**Table S6** Immunization schedule for generation of the anti-NltA1 polyclonal antibody.

| Date | Animal | Working task | Antiserum amount [mL] | Titer |
|---|---|---|---|---|
| 27 Jan 2026 | RB02263 | pre-immune serum/immunisation | 1.500 | |
| 10 Feb 2026 | RB02263 | boost | | |
| 24 Feb 2026 | RB02263 | boost | | |
| 10 Mar 2026 | RB02263 | bleeding/boost | 15.000 | X |
| 24 Mar 2026 | RB02263 | boost | | |
| 31 Mar 2026 | RB02263 | bleeding | 15.000 | |
| 14 Apr 2026 | RB02263 | boost | | |
| 28 Apr 2026 | RB02263 | final bleeding | 50.000 | |

#### 2. Supplementary figures

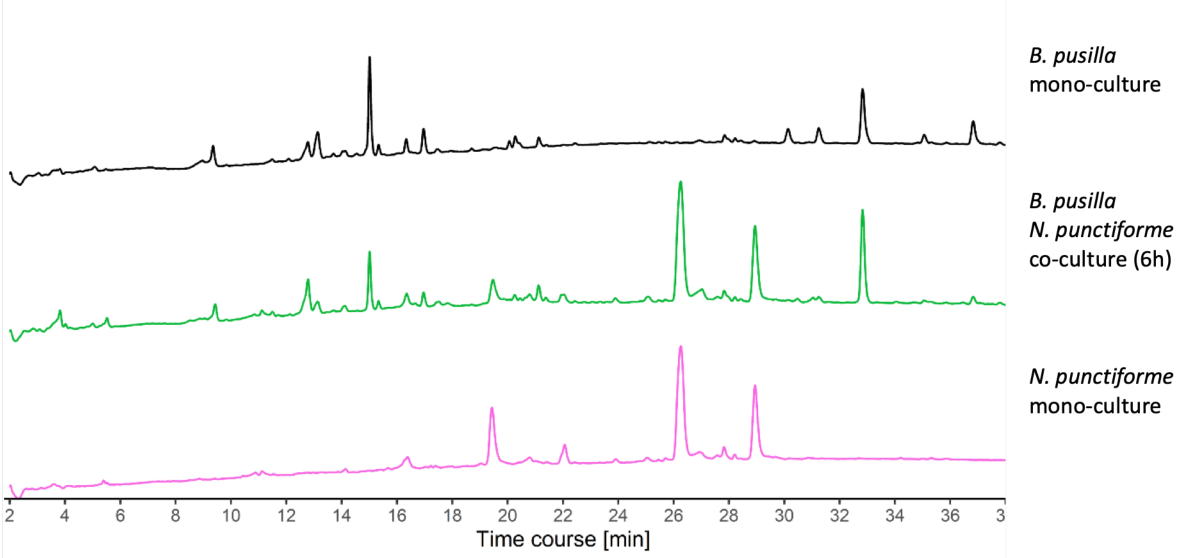


**Supplementary Figure 1. HPLC profiles of cell extracts of *B. pusilla* and *N. punctiforme* either grown in mono-culture of co-culture for 6 h.** Major peaks in the co-culture can either be traced back to *B. pusilla* or *N. punctiforme*, respectively. Targeted search for potential *ripp4* or *ripp3* products was unsuccessful.

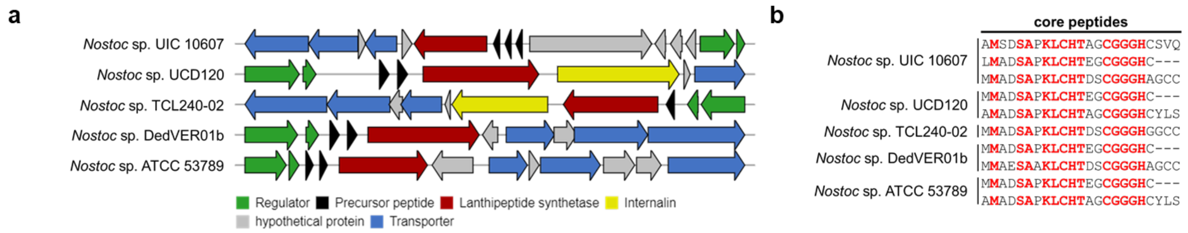


**Supplementary Figure 2. Representative BGCs closely related to the *nlt* BGC based on a BLASTP search of the precursor peptides, the lanthipeptide synthetase and the response regulators. a** The BGCs found in other *Nostoc* strains share the same overall architecture as the *nlt* BGC but harbor core peptides with a distinct conserved sequence motif, suggesting the biosynthesis of structurally different lanthipeptides. **b** Alignment of the corresponding core peptides. Highly conserved residues are highlighted in red.

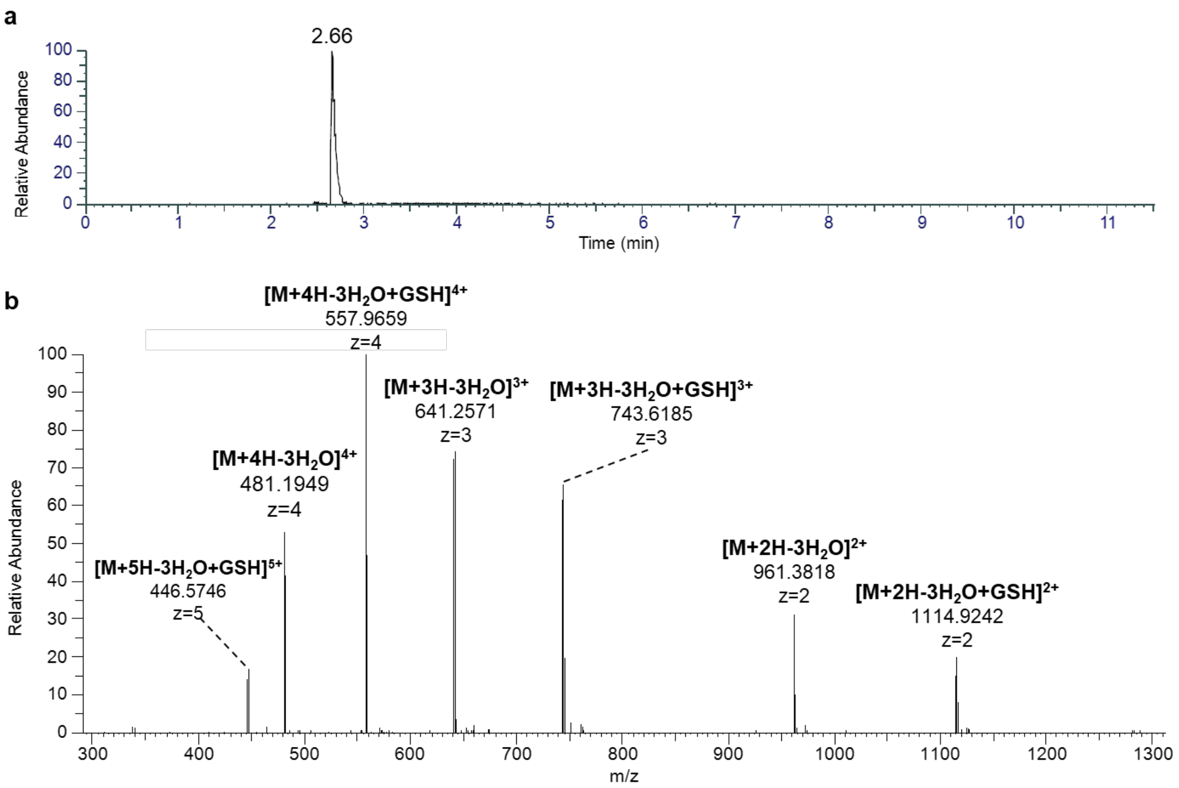


**Supplementary Figure 3. LC-MS analysis of LahT150-cleaved His$_6$-NltA1 co-expressed with NltM in *E. coli* identifies Nostolanthin A and a glutathionylated adduct. a** Extracted ion chromatogram (EIC) of the [M+3H-3$H_2O$]$^{3+}$ ion of Nostolanthin A (*m/z* 641.2571), showing a single peak at 2.66 min. **b** MS spectrum at the corresponding retention time, acquired on an Orbitrap mass spectrometer in positive ionization mode. Ion series corresponding to Nostolanthin A were observed as [M+2H-3$H_2O$]$^{2+}$ (*m/z* 961.3818), [M+3H-3$H_2O$]$^{3+}$ (*m/z* 641.2571), and [M+4H-3$H_2O$]$^{4+}$ (*m/z* 481.1949), consistent with loss of three molecules of water from the linear precursor and a monoisotopic mass of 1920.75 Da for the mature, fully cyclized peptide. A second series of ions with an additional mass shift of +307.0838 Da (corresponding to the mass of glutathione) was observed at [M+2H-3$H_2O$+GSH]$^{2+}$ (*m/z* 1114.9242), [M+3H-3$H_2O$+GSH]$^{3+}$ (*m/z* 743.6185), [M+4H-3$H_2O$+GSH]$^{4+}$ (*m/z* 557.9659), and [M+5H-3$H_2O$+GSH]$^{5+}$ (*m/z* 446.5746), consistent with covalent addition of glutathione to a dehydroalanine or dehydrobutyrine residue remaining after lanthionine ring formation.

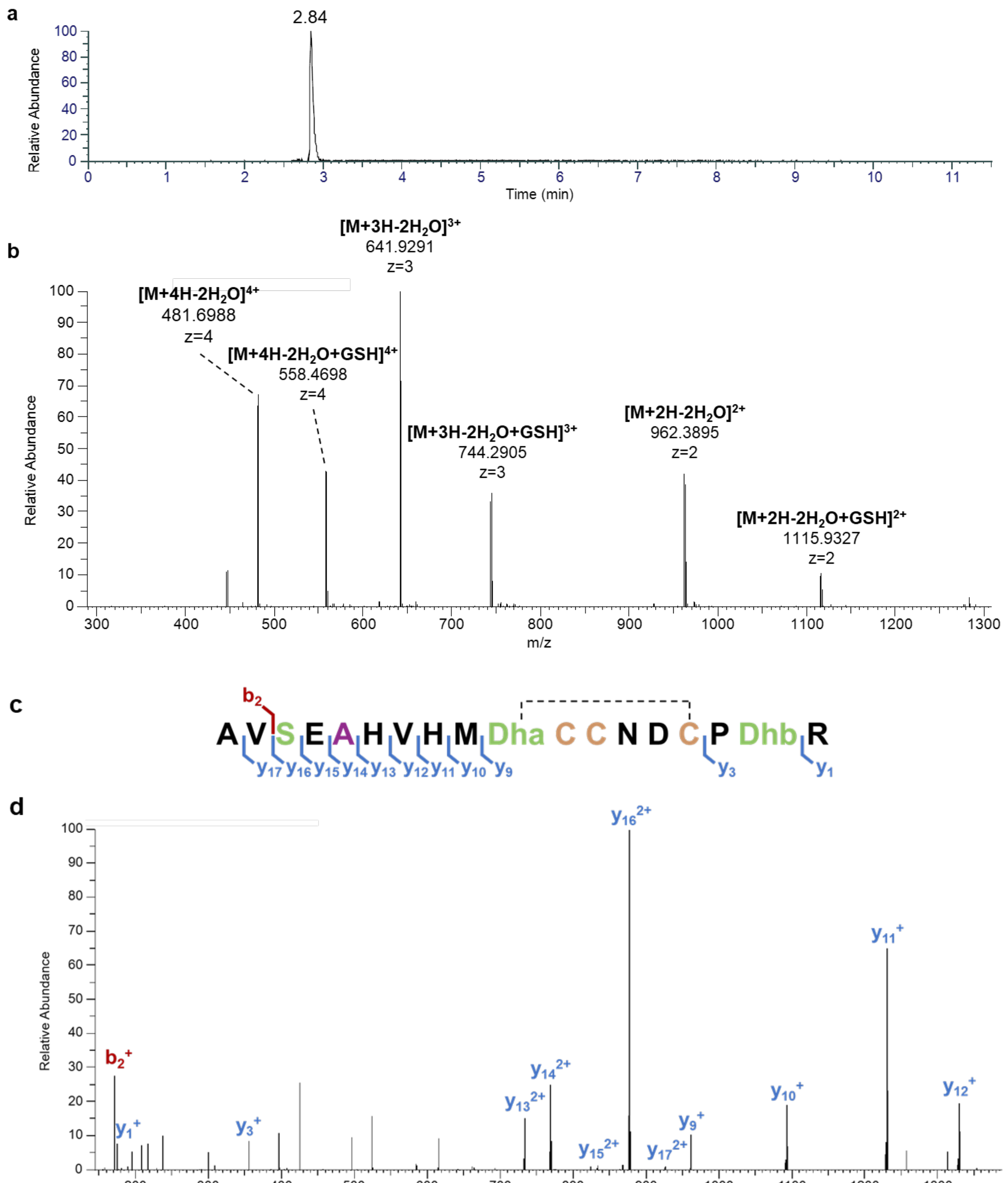


**e**

| ion | calculated mass [Da] | observed mass [Da] | error [ppm] |
|---|---|---|---|
| $b_2^+$ | 171.1128 | 171.1135 | +4.09 |
| $y_1^+$ | 175.1190 | 175.1200 | +5.71 |
| $y_3^+$ | 355.2088 | 355.2098 | +2.81 |
| $y_9^+$ | 962.3278 | 962.3209 | −7.17 |
| $y_{10}^+$ | 1093.3683 | 1093.3671 | −1.10 |
| $y_{11}^+$ | 1230.4272 | 1230.4288 | +1.30 |
| $y_{12}^+$ | 1329.4956 | 1329.5006 | +3.76 |
| $y_{13}^{2+}$ | 733.7809 | 733.7844 | +4.77 |
| $y_{14}^{2+}$ | 769.2995 | 769.2980 | −1.95 |
| $y_{15}^{2+}$ | 833.8208 | 833.8173 | −4.20 |
| $y_{16}^{2+}$ | 877.3368 | 877.3364 | −0.46 |
| $y_{17}^{2+}$ | 926.8710 | 926.8708 | −0.22 |

**Supplementary Figure 4. LC-MS and LC-MS/MS analysis of LahT150-cleaved His$_6$-NltA1-S5A co-expressed with NltM in *E. coli*. a** Extracted ion chromatogram (EIC) of the $[M+3H-2H_2O]^{3+}$ ion of nostolanthin-S5A (*m/z* 641.9291), showing a single peak at 2.84 min. **b** MS spectrum at the corresponding retention time, acquired on an Orbitrap mass spectrometer in positive ionization mode. Ion series corresponding to the S5A product were observed as $[M+2H-2H_2O]^{2+}$ (*m/z* 962.3895), $[M+3H-2H_2O]^{3+}$ (*m/z* 641.9291), and $[M+4H-2H_2O]^{4+}$ (*m/z* 481.6988), consistent with loss of only two molecules of water and a monoisotopic mass of 1958.79 Da, one fewer dehydration than observed for wild-type Nostolanthin A (Supplementary Fig. 3). This is consistent with the Ser5Ala substitution removing one of the three dehydroalanine residues. A second series of ions with an additional mass shift of +307.0838 Da (glutathione) was observed at $[M+2H-2H_2O+GSH]^{2+}$ (*m/z* 1115.9327), $[M+3H-2H_2O+GSH]^{3+}$ (*m/z* 744.2905), and $[M+4H-2H_2O+GSH]^{4+}$ (*m/z* 558.4698), consistent with covalent addition of glutathione to one of the two remaining unreacted dehydro-residues. **c** Sequence of the LahT150-cleaved peptide with observed b- and y-ions indicated above and below the corresponding fragment bonds, respectively. The dashed line indicates the proposed thioether connectivity of a lanthionine ring formed between Dha10 and Cys15. **d** MS/MS spectrum of the [*m/z* 641.93] precursor ion, with b- and y-type fragment ions labeled. **e** Calculated and observed monoisotopic masses and mass errors (ppm) for all assigned b- and y-ions shown in **c** and **d**.

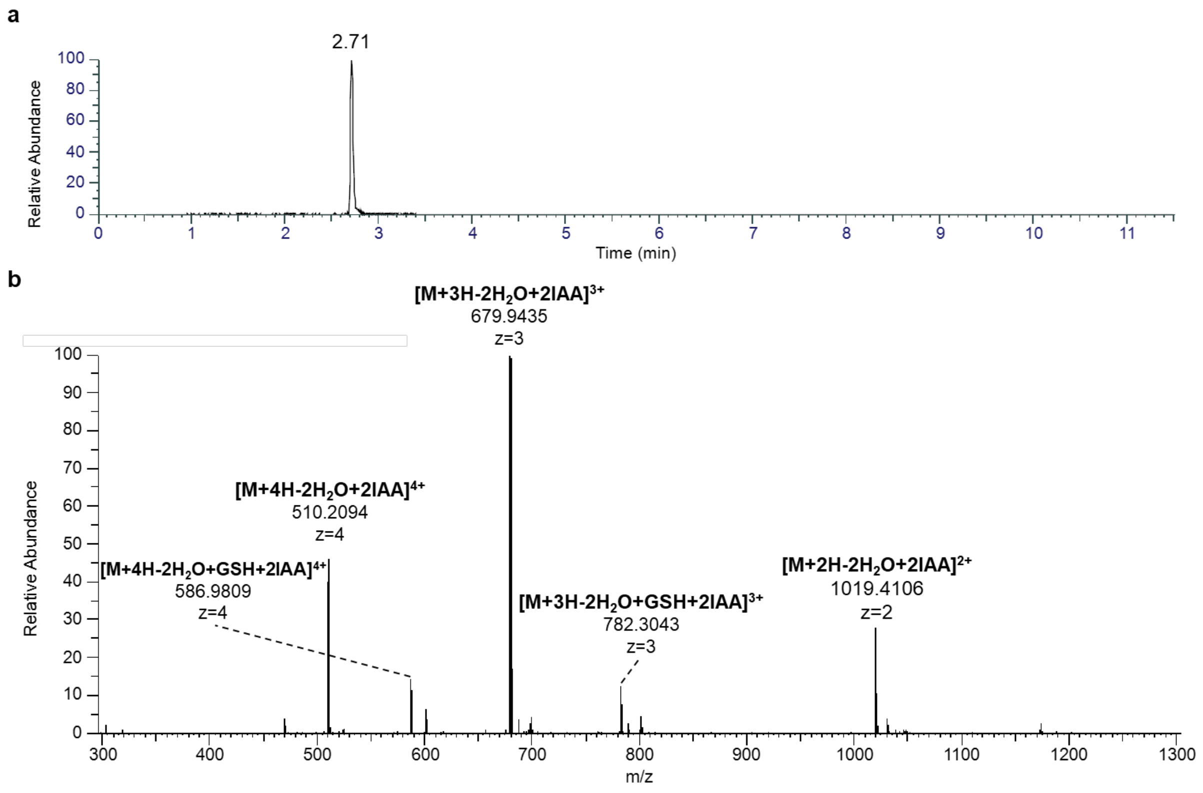


**Supplementary Figure 5. Iodoacetamide (IAA) assay of LahT150-cleaved His$_6$-NltA1-S5A co-expressed with NltM in *E. coli*. a** Extracted ion chromatogram (EIC) of the $[M+3H-2H_2O+2IAA]^{3+}$ ion of nostolanthin-S5A after IAA assay (*m/z* 679.9435), showing a single peak at 2.71 min. **b** MS1 spectrum showing dominant adducts of +2IAA across multiple charge states, indicating two free cysteine residues. Compared to wild-type nostolanthin, which carries one free cysteine, the S5A mutant retains an additional free cysteine, consistent with the loss of the lanthionine ring formed between Ser5 and its corresponding cysteine residue.

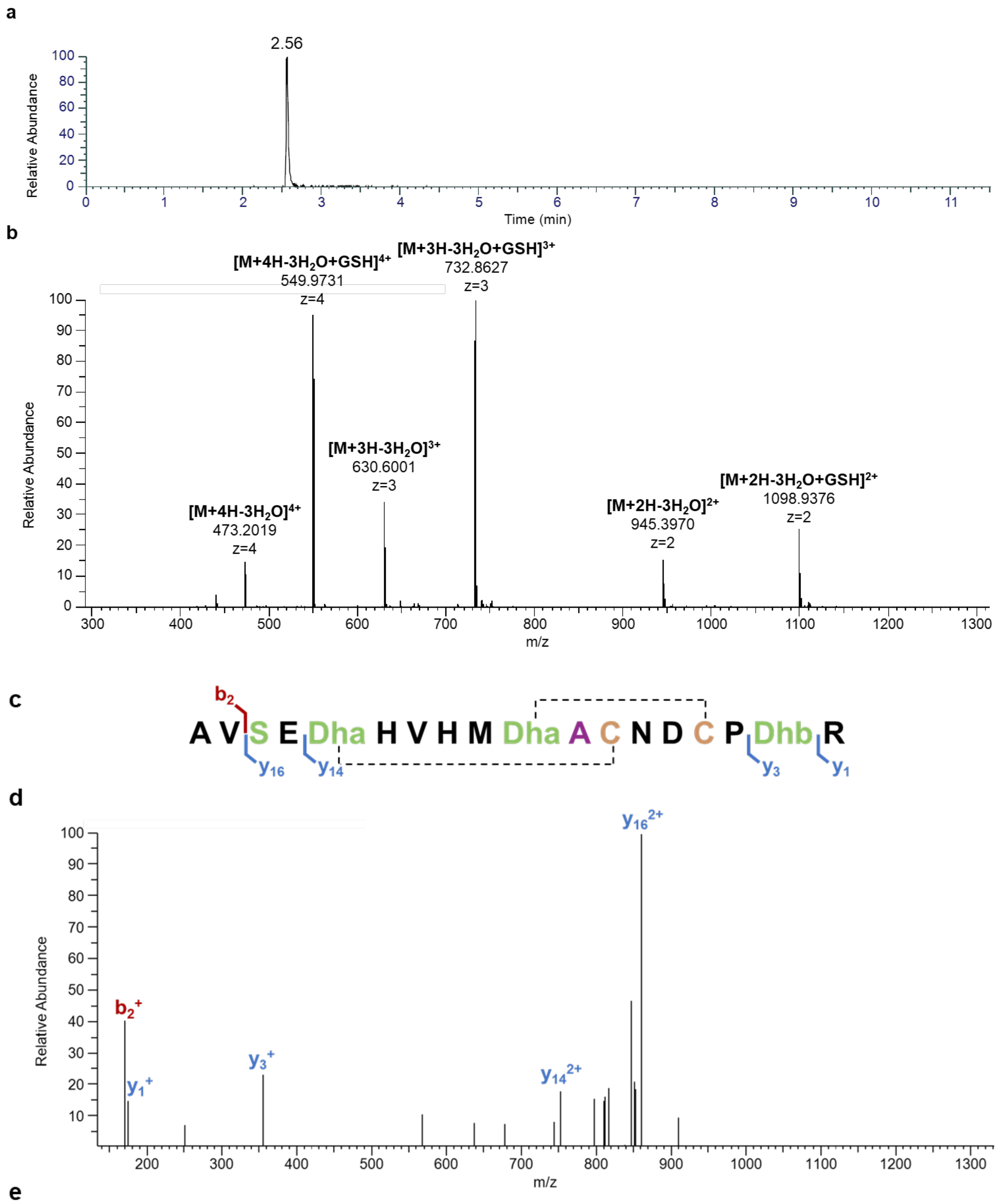


e

| ion | calculated mass [Da] | observed mass [Da] | error [ppm] |
|---|---|---|---|
| $b_2^+$ | 171.1128 | 171.1122 | −3.51 |
| $y_1^+$ | 175.1190 | 175.1190 | 0.00 |
| $y_3^+$ | 355.2088 | 355.2098 | +2.81 |
| $y_{14}^{2+}$ | 752.3056 | 752.3069 | +1.73 |
| $y_{16}^{2+}$ | 860.3429 | 860.3454 | +2.91 |

**Supplementary Figure 6. LC-MS and LC-MS/MS analysis of LahT150-cleaved His$_6$-NltA1-C11A co-expressed with NltM in *E. coli*. a** Extracted ion chromatogram (EIC) of the $[M+3H-3H_2O]^{3+}$ ion of nostolanthin-C11A (m/z 630.6001), showing a single peak at 2.56 min. **b** MS spectrum at the corresponding retention time, acquired on an Orbitrap mass spectrometer in positive ionization mode. Ion series corresponding to the C11A product were observed as $[M+2H-3H_2O]^{2+}$ (m/z 945.3970), $[M+3H-3H_2O]^{3+}$ (m/z 630.6001), and $[M+4H-3H_2O]^{4+}$ (m/z 473.2019), consistent with loss of three molecules of water and the same number of dehydrations as observed for wild-type Nostolanthin A, indicating that the Cys11Ala substitution does not abolish dehydration of any serine or threonine residue. A second series of ions with an additional mass shift of +307.0838 Da (glutathione) was observed at [M+2H-3$H_2$O+GSH]$^{2+}$ (m/z 1098.9376), [M+3H-3$H_2$O+GSH]$^{3+}$ (m/z 732.8627), and [M+4H-3$H_2$O+GSH]$^{4+}$ (m/z 549.9731), consistent with covalent addition of glutathione to one unreacted dehydro-residue. **c** Sequence of the LahT150-cleaved peptide with observed b- and y-ions indicated above and below the corresponding fragment bonds, respectively. The dashed line indicates the proposed thioether connectivity of a lanthionine ring. **d** MS/MS spectrum of the [m/z 630.93] precursor ion, with b- and y-type fragment ions labeled. **e** Calculated and observed monoisotopic masses and mass errors (ppm) for all assigned b- and y-ions shown in **c** and **d**.

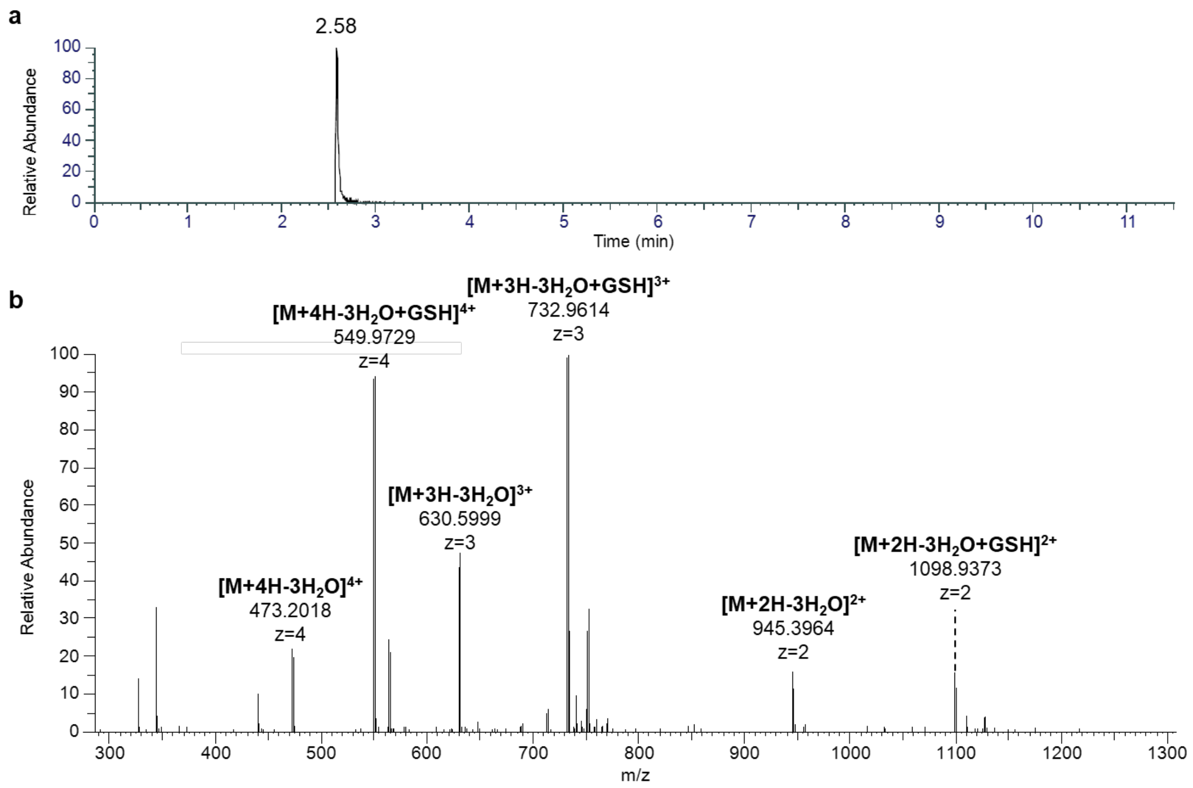


**Supplementary Figure 7. Iodoacetamide (IAA) assay of LahT150-cleaved His$_6$-NltA1-C11A co-expressed with NltM in *E. coli*. a** Extracted ion chromatogram (EIC) of the [M+3H-3$H_2$O]$^{3+}$ ion of nostolanthin-C11A product after IAA assay (m/z 630.5999), showing a single peak at 2.58 min. **b** MS1 spectrum showing no IAA adducts, indicating the absence of free cysteine residues. This demonstrates that both remaining cysteine residues in the C11A mutant are involved in thioether ring formation.

a

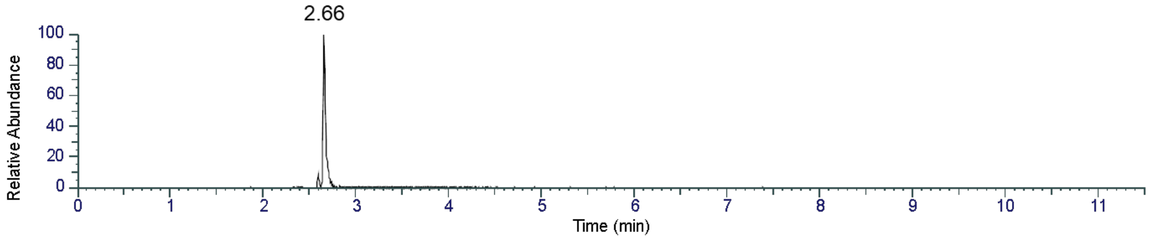


b

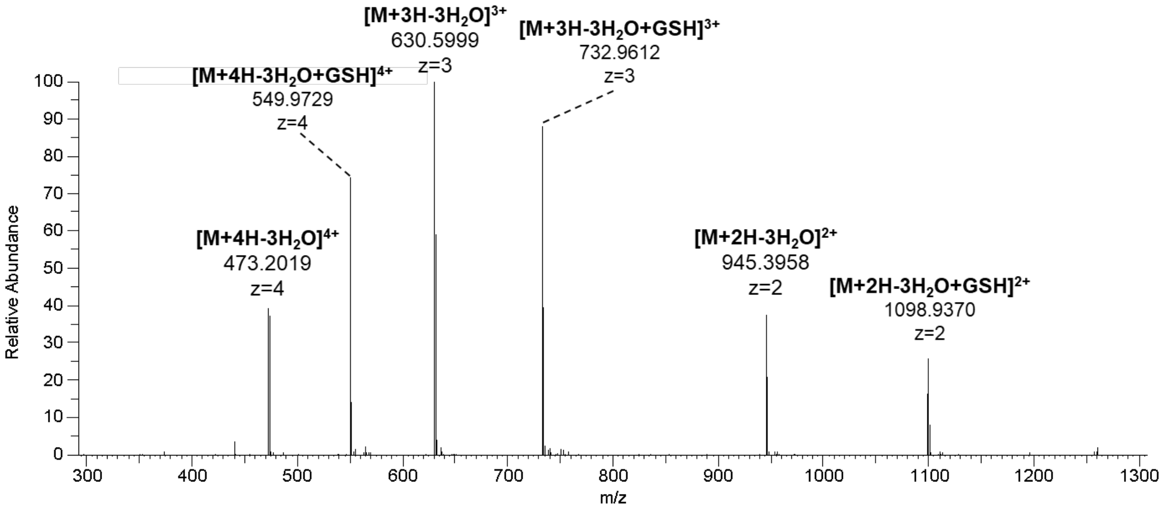


c

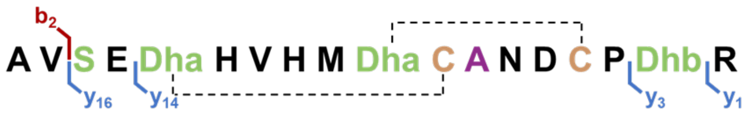


d

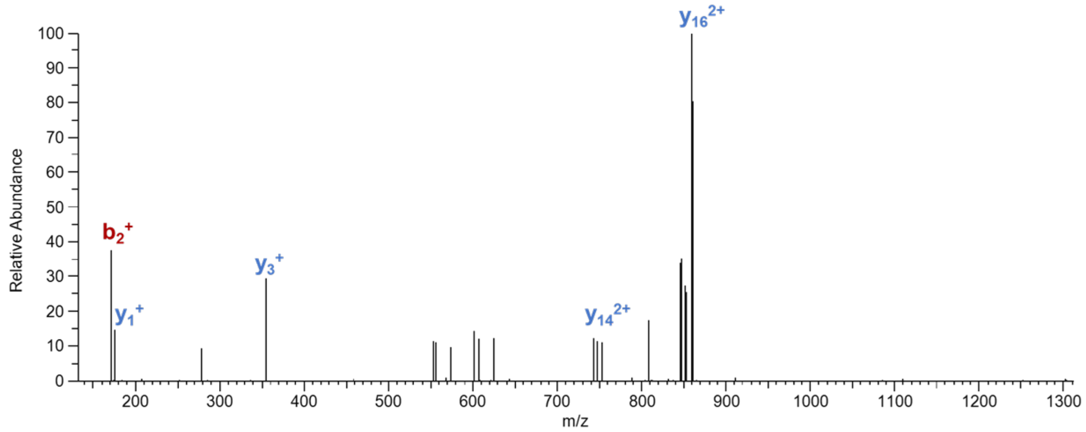


e

| ion | calculated mass [Da] | observed mass [Da] | error [ppm] |
|---|---|---|---|
| $b_2^+$ | 171.1128 | 171.1140 | +7.01 |
| $y_1^+$ | 175.1190 | 175.1191 | +0.57 |
| $y_3^+$ | 355.2088 | 355.2096 | +2.25 |
| $y_{14}^{2+}$ | 752.3056 | 752.3008 | −6.38 |
| $y_{16}^{2+}$ | 860.3429 | 860.3471 | +4.88 |

**Supplementary Figure 8. LC-MS and LC-MS/MS analysis of LahT150-cleaved His$_6$-NltA1-C12A co-expressed with NltM in *E. coli*. a** Extracted ion chromatogram (EIC) of the $[M+3H-3H_2O]^{3+}$ ion of nostolanthin-C12A (m/z 630.5999), showing a single peak at 2.66 min. **b** MS spectrum at the corresponding retention time, acquired on an Orbitrap mass spectrometer in positive ionization mode. Ion series corresponding to the C12A product were observed as $[M+2H-3H_2O]^{2+}$ (m/z 945.3958), $[M+3H-3H_2O]^{3+}$ (m/z 630.5999), and $[M+4H-3H_2O]^{4+}$ (m/z 473.2019), consistent with loss of three molecules of water and the same number of dehydrations as observed for wild-type Nostolanthin A, indicating that the Cys12Ala substitution does not abolish dehydration of any serine or threonine residue. A second series of ions with an additional mass shift of +307.0838 Da (glutathione) was observed at [M+2H-

3$H_2$O+GSH]$^{2+}$ (m/z 1098.9370), [M+3H-3$H_2$O+GSH]$^{3+}$ (m/z 732.9612), and [M+4H-3$H_2$O+GSH]$^{4+}$ (m/z 549.9729), consistent with covalent addition of glutathione to one unreacted dehydro-residue. **c** Sequence of the LahT150-cleaved peptide with observed b- and y-ions indicated above and below the corresponding fragment bonds, respectively. The dashed line indicates the proposed thioether connectivity of a lanthionine ring. **d** MS/MS spectrum of the [m/z 630.93] precursor ion, with b- and y-type fragment ions labeled. **e** Calculated and observed monoisotopic masses and mass errors (ppm) for all assigned b- and y-ions shown in **c** and **d**.

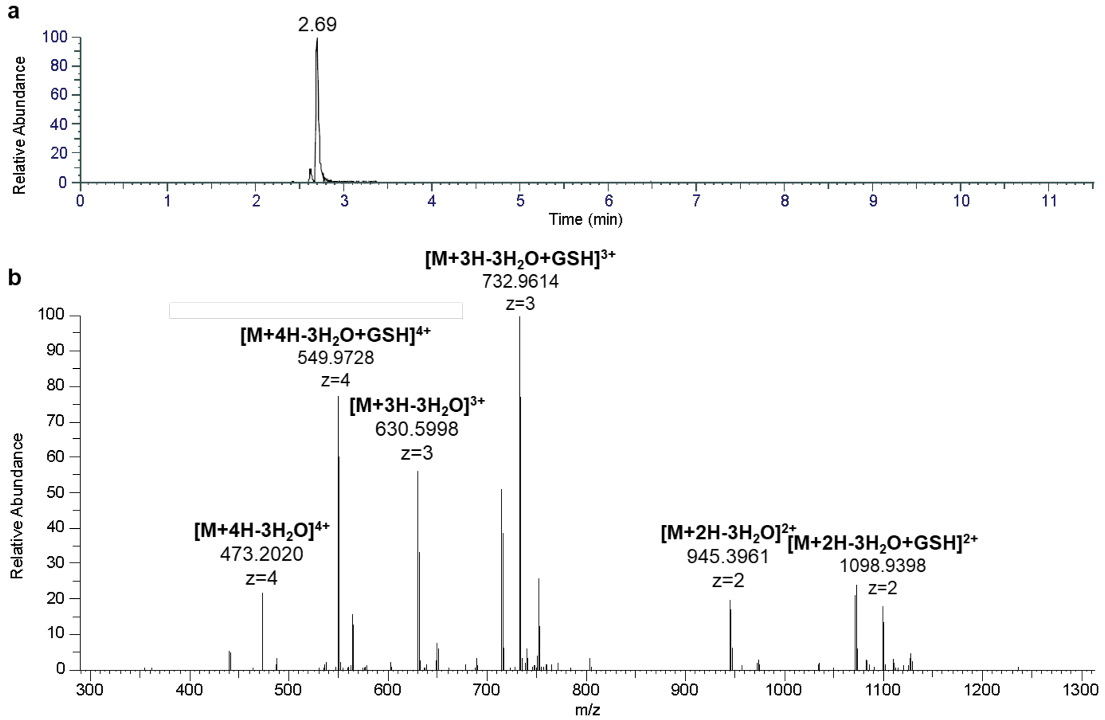


**Supplementary Figure 9. Iodoacetamide (IAA) assay of LahT150-cleaved His$_6$-NltA1-C12A co-expressed with NltM in *E. coli*. a** Extracted ion chromatogram (EIC) of the [M+3H-3$H_2$O]$^{3+}$ ion of nostolanthin-C12A product after IAA assay (m/z 630.5998), showing a single peak at 2.69 min. **b** MS1 spectrum showing no IAA adducts, indicating the absence of free cysteine residues. This demonstrates that both remaining cysteine residues in the C12A mutant are involved in thioether ring formation.

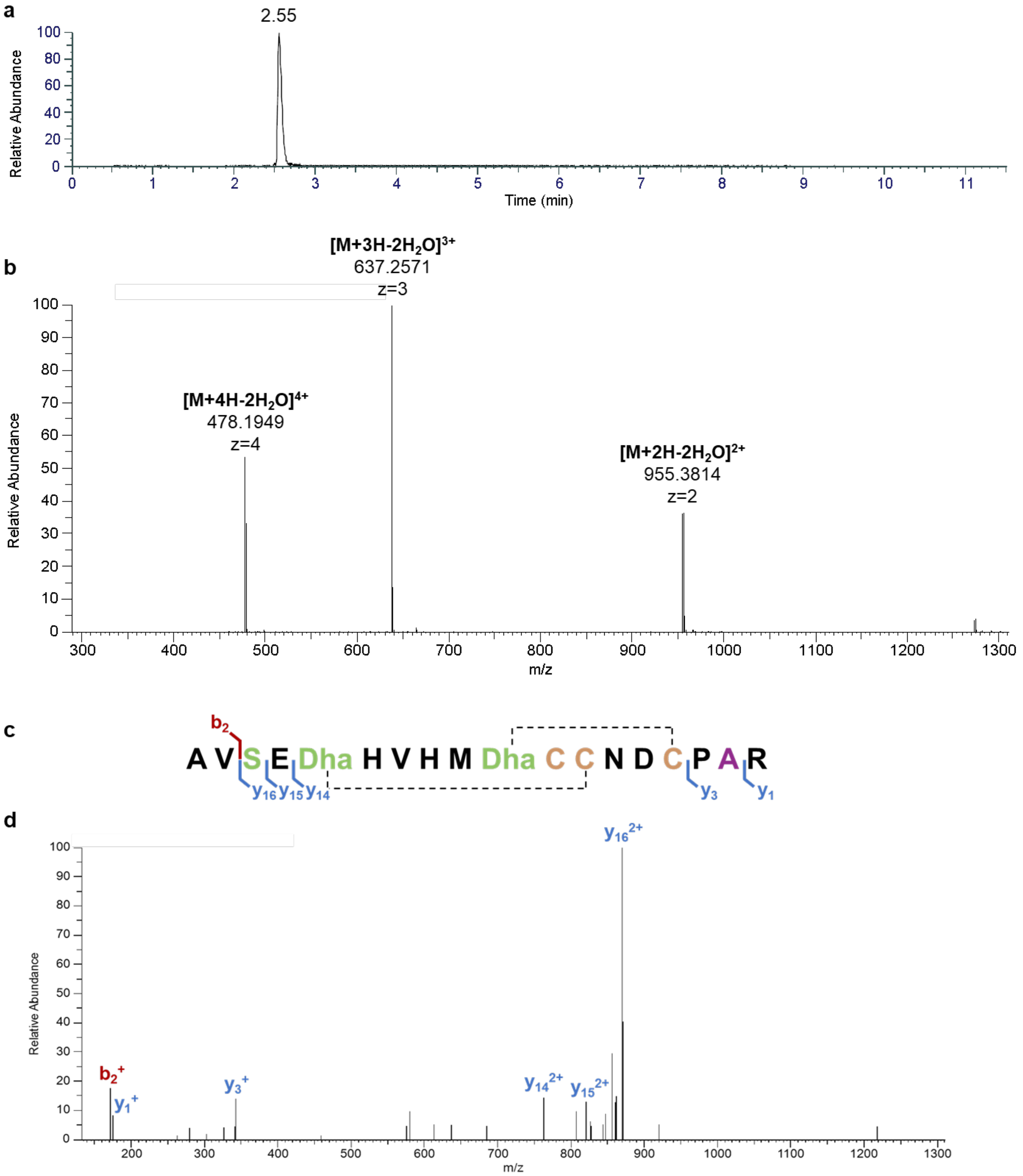


**e**

| ion | calculated mass [Da] | observed mass [Da] | error [ppm] |
|---|---|---|---|
| $b_2^+$ | 171.1128 | 171.1134 | +3.51 |
| $y_1^+$ | 175.1190 | 175.1192 | +1.14 |
| $y_3^+$ | 343.2089 | 343.2086 | −0.87 |
| $y_{14}^{2+}$ | 762.2916 | 762.2920 | +0.52 |
| $y_{15}^{2+}$ | 826.8129 | 826.8171 | +5.08 |
| $y_{16}^{2+}$ | 870.3290 | 870.3285 | −0.57 |

**Supplementary Figure 10. LC-MS and LC-MS/MS analysis of LahT150-cleaved His$_6$-NltA1-T17A co-expressed with NltM in *E. coli*. a** Extracted ion chromatogram (EIC) of the $[M+3H-2H_2O]^{3+}$ ion of nostolanthin-T17A (m/z 637.2571), showing a single peak at 2.55 min. **b** MS spectrum at the corresponding retention time, acquired on an Orbitrap mass spectrometer in positive ionization mode. Ion series corresponding to the T17A product were observed as $[M+2H-2H_2O]^{2+}$ (m/z 955.3814), $[M+3H-2H_2O]^{3+}$ (m/z 637.2571), and $[M+4H-2H_2O]^{4+}$ (m/z 478.1949), consistent with loss of only two molecules of water and one fewer dehydration than observed for wild-type Nostolanthin A. This is consistent with the Thr17Ala substitution removing the dehydrobutyrine residue at position 17. Notably, no glutathione adducts were detected, in contrast to wild-type and other mutants, indicating the absence of any remaining unreacted dehydro-residues and confirming that Dhb17 is the free dehydro amino acid that is glutathionylated in the wild-type peptide. **c** Sequence of the LahT150-cleaved peptide with observed b- and y-ions indicated above and below the corresponding fragment bonds, respectively. The dashed line indicates the proposed thioether connectivity of the two lanthionine rings. **d** MS/MS spectrum of the [m/z 637.25] precursor ion, with b- and y-type fragment ions labeled. **e** Calculated and observed monoisotopic masses and mass errors (ppm) for all assigned b- and y-ions shown in **c** and **d**.

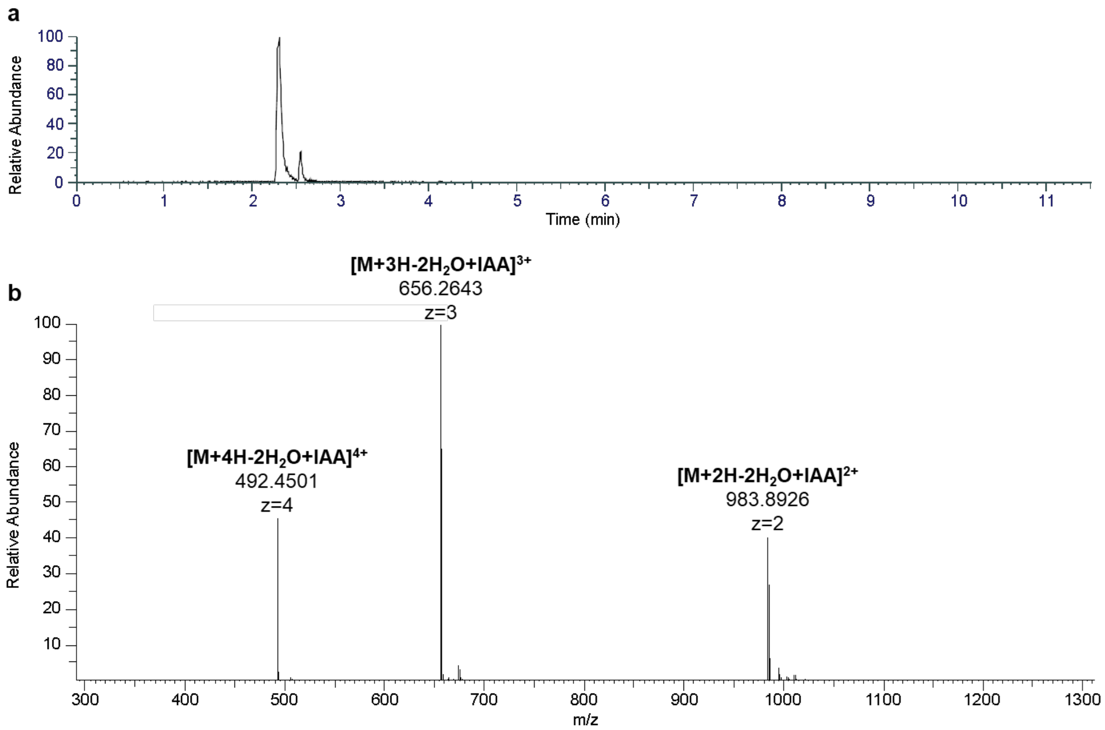


**Supplementary Figure 11. Iodoacetamide (IAA) assay of LahT150-cleaved His$_6$-NltA1-T17A co-expressed with NltM in *E. coli*. a** Extracted ion chromatogram (EIC) of the [M+3H-2H$_2$O+IAA]$^{3+}$ ion of nostolanthin-T17A product after IAA assay (m/z 656.2643), showing two peaks at 2.30 and 2.55 min. **b** MS1 spectrum showing dominant adducts of +1IAA across multiple charge states, as observed at [M+2H-2H$_2$O+IAA]$^{2+}$ (m/z 983.8926), [M+3H-2H$_2$O+IAA]$^{3+}$ (m/z 656.2643), and [M+4H-2H$_2$O+IAA]$^{4+}$ (m/z 492.4501), indicating one free cysteine residue. This is consistent with the Thr17Ala substitution confirming that Thr17 in wild-type corresponds to the free dehydro amino acid that is glutathionylated, while one cysteine remains uninvolved in ring formation, mirroring the free cysteine observed in wild-type Nostolanthin A.

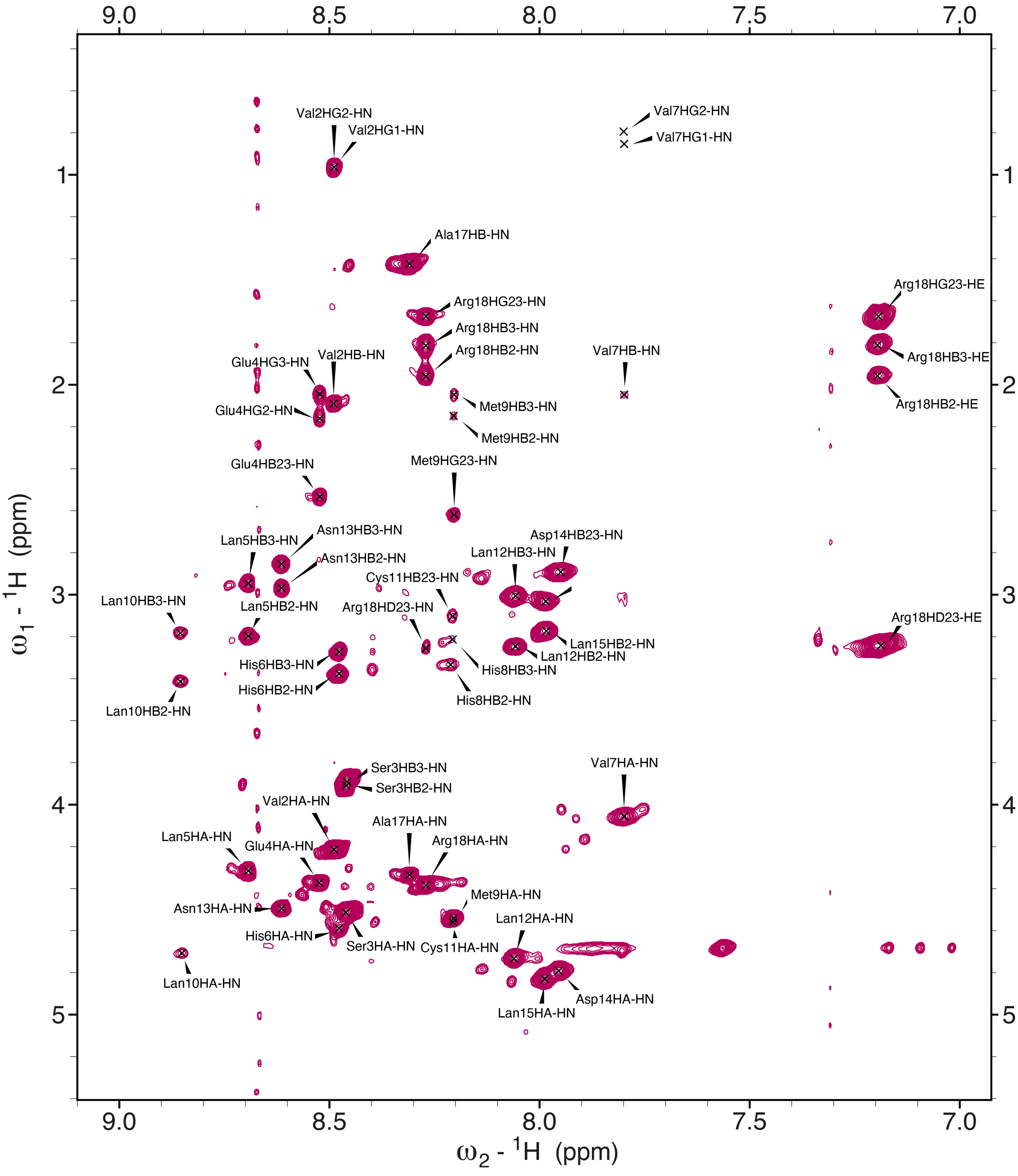


**Supplementary Figure 12. Amide region of the $^{1}H$-$^{1}H$ TOCSY spectrum of nostolanthin-T17A recorded in H2O at 308 K.** Cross-peaks are labeled with resonance assignments. Notably, the amide resonances of His8, Met9 and Cys11 overlap at 8.2 ppm.

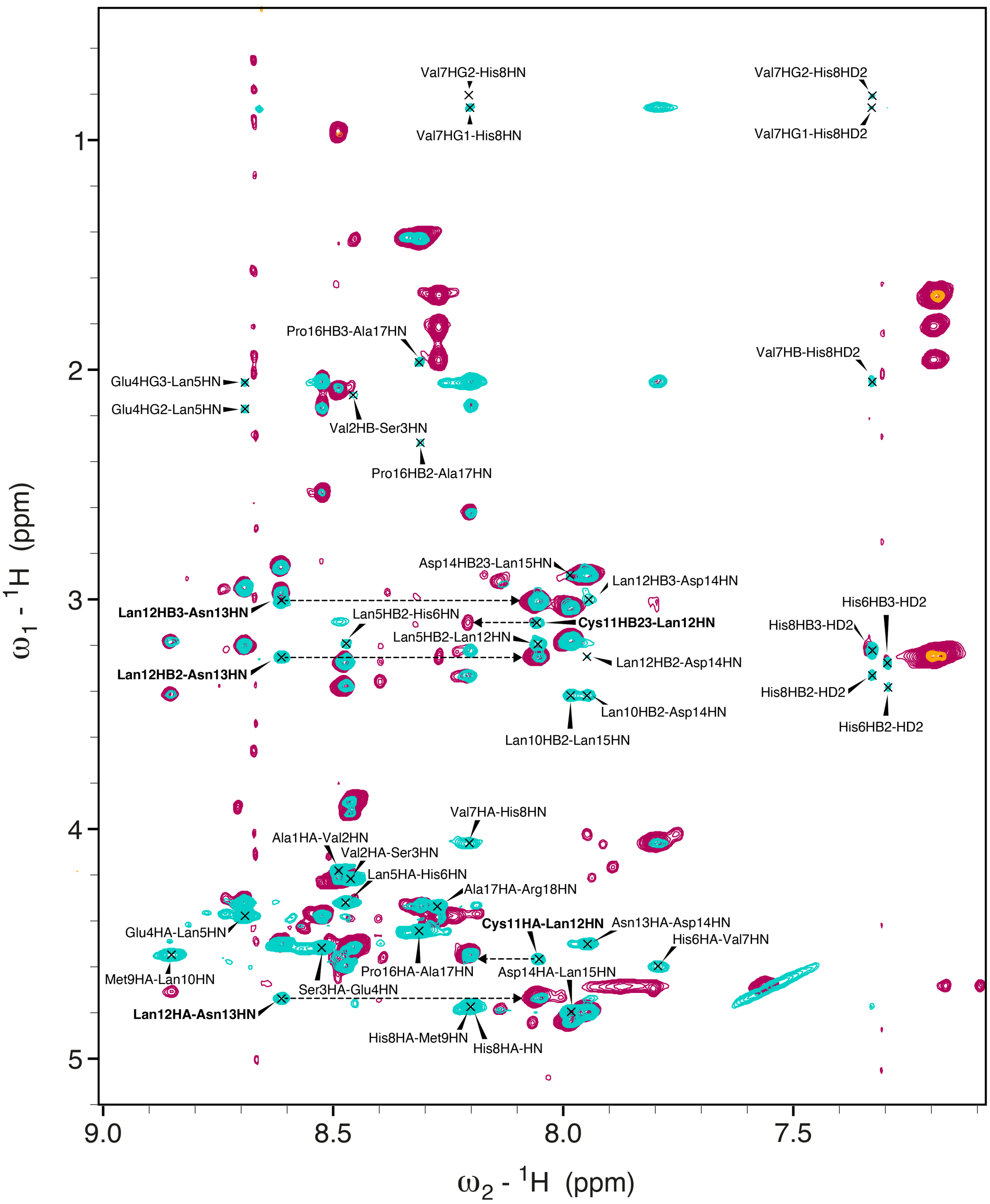


**Supplementary Figure 13. Overlay of the amide regions of $^1$H-$^1$H TOCSY (magenta) and $^1$H-$^1$H NOESY (cyan) spectra of nostolanthin-T17A recorded in $H_2O$ at 308 K.** Only inter-residual cross-peaks are labeled (compare with intra-residual TOCSY correlations in Figure S12). Sequential key correlations between residues Cys11, Lan12 and Asn13 are highlighted in bold and with dashed arrows.

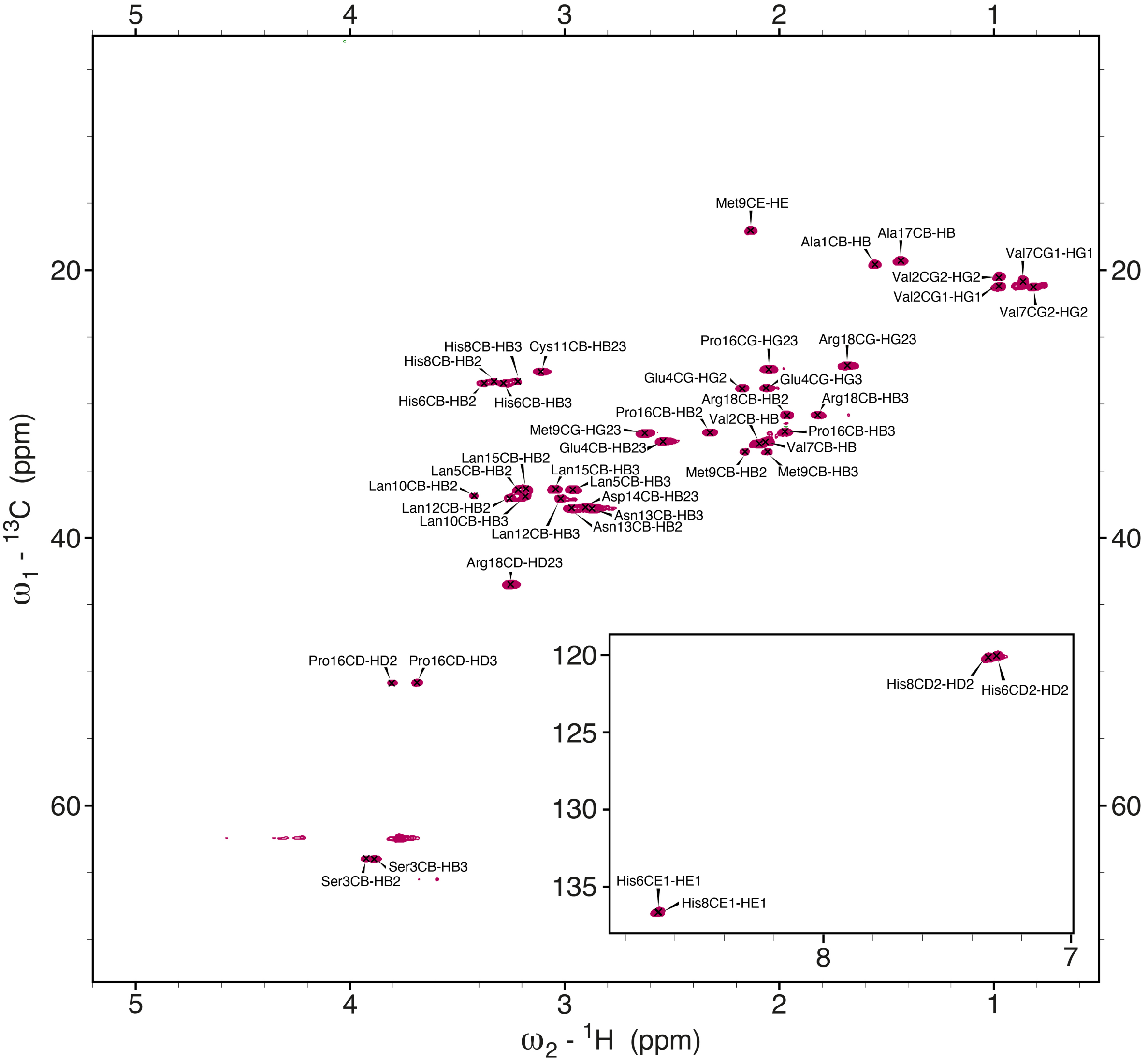


**Supplementary Figure 14. $^1$H-$^{13}$C HSQC spectrum of nostolanthin-T17A recorded in $H_2O$ at 308 K.** Cross-peaks are labeled with resonance assignments. Resonances of CHα groups were obscured by the water signal at 4.8 ppm and thus not observed. The imidazole region of His6 and His8 is depicted in the inset.

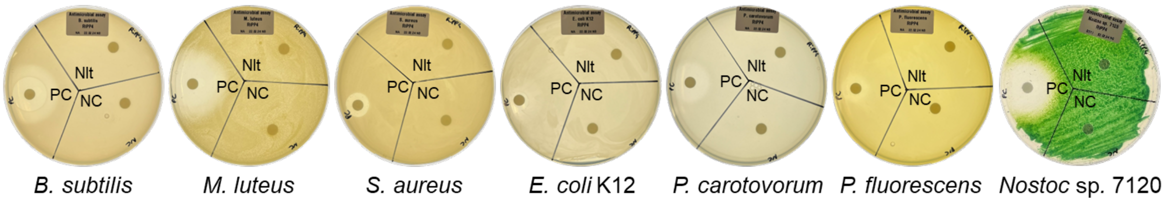


**Supplementary Figure 15. Bioactivity assay with nostolanthin A.** 10 µg of nostolanthin A (Nlt) against the Gram-positive bacteria *B. subtilis*, *M. luteus*, *S. aureus* and the Gram-negative bacteria *E. coli* K12, *P. carotovorum*, *P. fluorescens* and *Nostoc* sp. 7120. Water served as negative control (NC), 20 µg streptomycin served as positive control (PC).

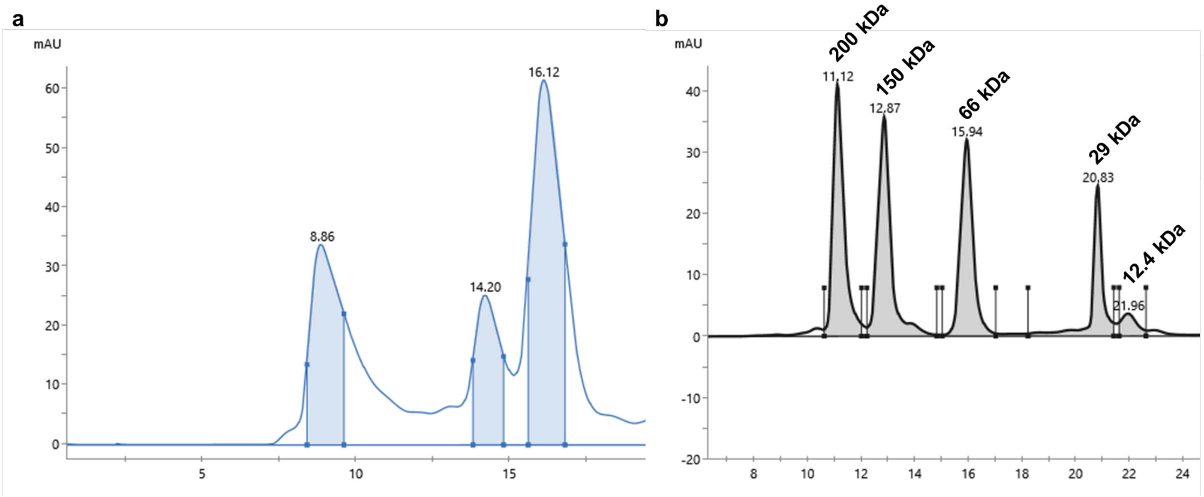


**Supplementary Figure 16. Analytical size-exclusion chromatography (SEC) analysis of NltA1. a** SEC chromatogram of the NltA1 precursor peptide co-expressed with NltM in *E. coli*, purified by affinity chromatography, and subsequently incubated in pH 10.6 buffer for 48 h. A high-molecular-weight species could be enriched at an elution volume of $V_e$: 8.86 mL. **b** SEC calibration curve generated using the Gel Filtration Molecular Weight Markers Kit (MWGF200, Sigma-Aldrich; molecular weight range 12,000–200,000 Da). Standards used: cytochrome c (12.4 kDa), carbonic anhydrase (29 kDa), bovine serum albumin (66 kDa), alcohol dehydrogenase (150 kDa), and β-amylase (200 kDa).

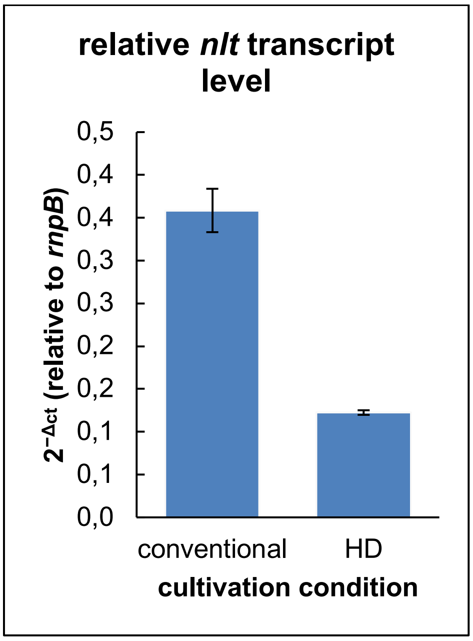


**Supplementary Figure 17. RT-qPCR of the *nlt* BGC in *Nostoc* sp. KVJ2-*nlt*.** The relative transcript levels are normalized to *rnpB*.

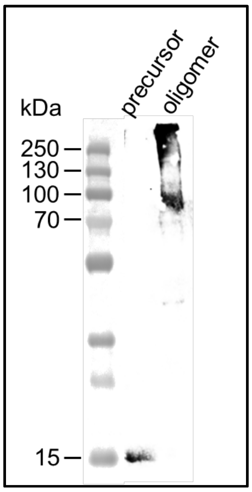


**Supplementary Figure 18. Western blot to test the polyclonal antibody anti-NltA1.** The antibody recognizes both, the nostolanthin precursor peptide monomer and oligomer, both purified from *E. coli* after co-expression with NltM and separated using SEC.
